# cloneXplorer: A high-throughput clone discovery platform based on conical microwell arrays

**DOI:** 10.64898/2026.01.16.699323

**Authors:** Guido K. Stadler, Eugene Tkachenko, Oscar Neri, Michael Zakharov, Orr Zohar, Debbie X. Deng, Kitt D. Paraiso, Hajar Rajaei, Shaun Steele, Xiling Shen, Alex Chenchik, Benjamin B. Yellen

## Abstract

Antigen-reactive T cell populations are of great value for studying immune recognition but tedious to generate by limiting dilution or cloning. Here, we develop a streamlined approach to generate antigen-reactive T cell clones directly from peripheral blood using the cloneXplorer, a live-cell analysis and clone isolation platform based on conical microwell arrays. This platform continuously monitors cell proliferation, cytokine secretion, and surface markers in up to 100,000 single cell co-cultures, enabling the identification of rare, functionally defined T cells, which can be recovered for clonal expansion or sequence analysis. We benchmark the platform by performing several key demonstrations. First, we show that this platform can efficiently generate monoclonal cell populations from cell lines and human primary T cells. Next, we demonstrate that antigen-specificity can be identified at single cell resolution using Jurkat cells expressing NFAT-GFP, CD8, and a T cell receptor (TCR) co-cultured with K562 antigen presenting cells (APC) expressing a peptide library. Thereafter, we show that immune activation in mouse and human primary samples can be monitored by time lapse analysis of Interferon gamma (IFN-γ) secretion in individual microwell co-cultures using a fluorescent sandwich assay. Finally, we combine these capabilities in a proof-of-concept demonstration, which uses IFN-γ secretion and the presence of CD8 surface markers as hierarchical gates to isolate and expand antigen-reactive T cells from human peripheral blood. We verify the reactivity of the expanded cells both using tetramer staining and functional re-challenge assays. We also re-express the identified TCRs in a Jurkat reporter cell line to validate the specificity of the matched TCR/antigen pairing in an artificial system. Together, these results showcase potential applications of the cloneXplorer platform in cell line development, and in screening and validating immune receptor interactions with specific antigens.

## INTRODUCTION

Single cell assays can quantify critical information about individual cells that would otherwise be lost in the analysis of bulk cell populations. For example, the ability to determine both the alpha and beta chain sequences of the T cell receptor (TCR), or the heavy and light chains of the B cell receptor, necessitates measurement of two distinct molecules originating from the same cell^1,2^. Many approaches have been developed to match TCRs with their antigen target using single cell TCR sequencing (scTCR-seq) in combination with methods to enrich the population based on multimer binding^3,4^, activation marker expression^1,5^, or functional readouts, such as cytokine secretion^6–8^ or target cell killing^9^. The destructive nature of TCR-seq usually requires additional steps for functional verification of antigen-TCR pairing, including cloning and expressing the TCR in primary T cells or reporter cell lines. To overcome this limitation, we developed an automated platform to identify and expand the antigen-specific T cell clones with a functional screen, which allowed us to use the same single T cells both for TCR sequence identification and as the precursor to generate a monoclonal cell line, which are later used in a re-challenge assay to confirm specificity of the TCR-antigen interaction. This capability dramatically simplifies antigen validation and TCR discovery workflows, and also streamlines the development and testing of cancer vaccines and cell-based therapies by obviating the need for molecular cloning steps when establishing monoclonal T cell populations.

Here, we showcase a potential application of the cloneXplorer^TM^ in establishing a streamlined antigen validation workflow, in which T cells with desired antigen reactivity are discovered and expanded into functionally defined monoclonal T cell populations. To achieve this goal, we built a fully automated cell discovery platform based on an array of 380,000 conically shaped microwells and a live cell imaging instrument that automatically scans and analyzes each microwell over time to pinpoint the best cells for isolation. We assess basic performance metrics of the platform by measuring cell loading statistics, single cell proliferation rates, and by demonstrating that desired cells can be retrieved with a computer vision guided syringe tip and transferred to the deposition plate.

We benchmark the performance of the cloneXplorer by demonstrating multiple biological workflows. First, we establish a cell line development workflow, where we show that cloneXplorer can be used to generate monoclonal cell populations from cell lines and human T cells with efficiency as high as 95%. Next, we develop an antigen screening workflow based on monitoring fluorescent reporter activation in single Jurkat T cells co-cultured with a peptide construct library expressed in K562 antigen presenting cells (APC), where we use sequencing to identify the singly presented antigen in the identified microwells. Third, we establish a robust cytokine detection workflow by performing time-resolved measurement of Interferon gamma (IFN-γ) secretion from mouse T cells and APCs co-cultured in microwells, allowing this functional information to be used in combination with surface marker expression and cell proliferation measurements to identify the best clones for isolation and expansion. Finally, we developed a pipeline for identifying specific TCR-antigen interactions using these expanded cell populations from donor blood both for TCR sequencing and TCR-antigen affinity measurements in a functional rechallenge assay. While other approaches can identify TCRs specific to a target antigen using droplets or microfluidic devices, we are unaware of any approach that can both identify TCR sequences and validate function in a re-challenge assay using materials derived from the same single cell. Taken together, the cloneXplorer platform enables the discovery and quantification of antigen-reactive T cells, the screening of complex antigen and/or TCR libraries for functional interactions and supports workflows for developing cancer vaccines and T cell-based therapies.

## RESULTS

### Working concept of the cloneXplorer

We set out to develop an instrument and consumable to screen rare cells with a specific phenotypic function and isolate them for downstream analyses. The consumable consists of a microwell array bonded to a 96 well plate frame (**Fig.1a**, **S1**), which is designed in standard Society for Laboratory Automation and Screening (SLAS) footprint to make it compatible with most imagers and liquid handlers. The unique geometry of these microwells make them ideally suited for assays involving single cells or defined co-cultures. First, the conical shape with intersecting walls enables all seeded cells to be collected at the bottom of the microwells by centrifugation. Second, the high aspect ratio (height / width > 2) provides enough space to grow more than 100 cells per microwell during cell expansion.

High aspect ratio also makes it possible to perform liquid exchanges without disturbing the cells inside the microwells. Our manufacturing process can produce microwells with diameters as small as 7 µm and as big as 250 µm for specialized applications, such as analysis of bacteria, yeast, and organoids (**Fig. S2**). The materials used to manufacture the arrays were selected for compatibility with fluorescent imaging and to reduce the adsorption of proteins and small molecules, except where desired.

To automate cell assays, we built the cloneXplorer, which is an instrument that combines a cell culture incubator, an imager, and a cell picker (**Fig.1b, S3**) into an integrated functional cell discovery and isolation platform. The incubator employs heating elements in the enclosure walls to maintain uniform temperature, while the stage-top incubator controls humidity, O_2_ and CO_2_ (**Fig.1b E, C**). The imager (**Fig.1b F-K**) consists of an inverted epi-fluorescent microscope capable of high-speed image acquisition (<40 min for full plate scan in 4 fluorescent channels at 100 ms exposures plus brightfield with autofocusing at every field of view) at a resolution of 0.5 µm/pixel.

The diagram in **Fig.1c** shows a typical workflow for isolating proliferative single cells. First, the microwell array is primed with cell culture medium. Next, cells are dispensed into each well and they sediment into the microwells by gravity. The plate is then loaded into the cloneXplorer for live cell imaging. Due to the open microwell format, the ∼1 nL volume of the microwell shares media with the larger well volume (∼0.3 mL), so that soluble molecules, such as nutrients and metabolites, can diffuse in and out of microwells, allowing cells to be grown and imaged for days without feeding. Real time image analysis is used to select microwells based on various phenotypic measurements. For example, **Fig.1c** depicts the identification of a clone that expands from a single cell. This clone can be retrieved by the picker, which consists of a syringe mounted on a XYZ stage (**Fig.1b O-P**). The conical “funnel” shape of the microwell guides the picker tip into an optimal picking position without the need for fine tip control and is essential for fully automating the picking process. We use a 36-gauge steel needle that fits perfectly into the microwell opening but cannot reach the bottom, ensuring that cells are not damaged during recovery. The conical shape is also ideal for establishing a pressure seal between the picker tip and the rim of the microwell due to the constriction fit made upon contact. The picking procedure involves moving the syringe barrel to the 1μL position without breaking contact with the microwell, which builds up negative pressure. When the tip breaks contact, it instantaneously generates hydrodynamic disturbance that produces internal convection leading to withdrawal of the microwell contents. Retrieved cells are transferred into a specific well of the 96-well deposition plate (**Fig.1b Q**) after which the tip is rinsed in the wash plate (**Fig.1b R**) and readied for the next pick. Laminar flow from the HEPA filter located at the ceiling of the instrument maintains particle-free environment during the retrieval process (**Fig.1b N**) and rapidly cools the instrument to room temperature to reduce evaporation during picking. Using this approach, we consistently achieved >95% efficiency in retrieving both for 7 µm polymer beads and NALM6-GFP cells from the microwells (**Fig. 1d, Movies S1-2**).

**Fig. 1.**
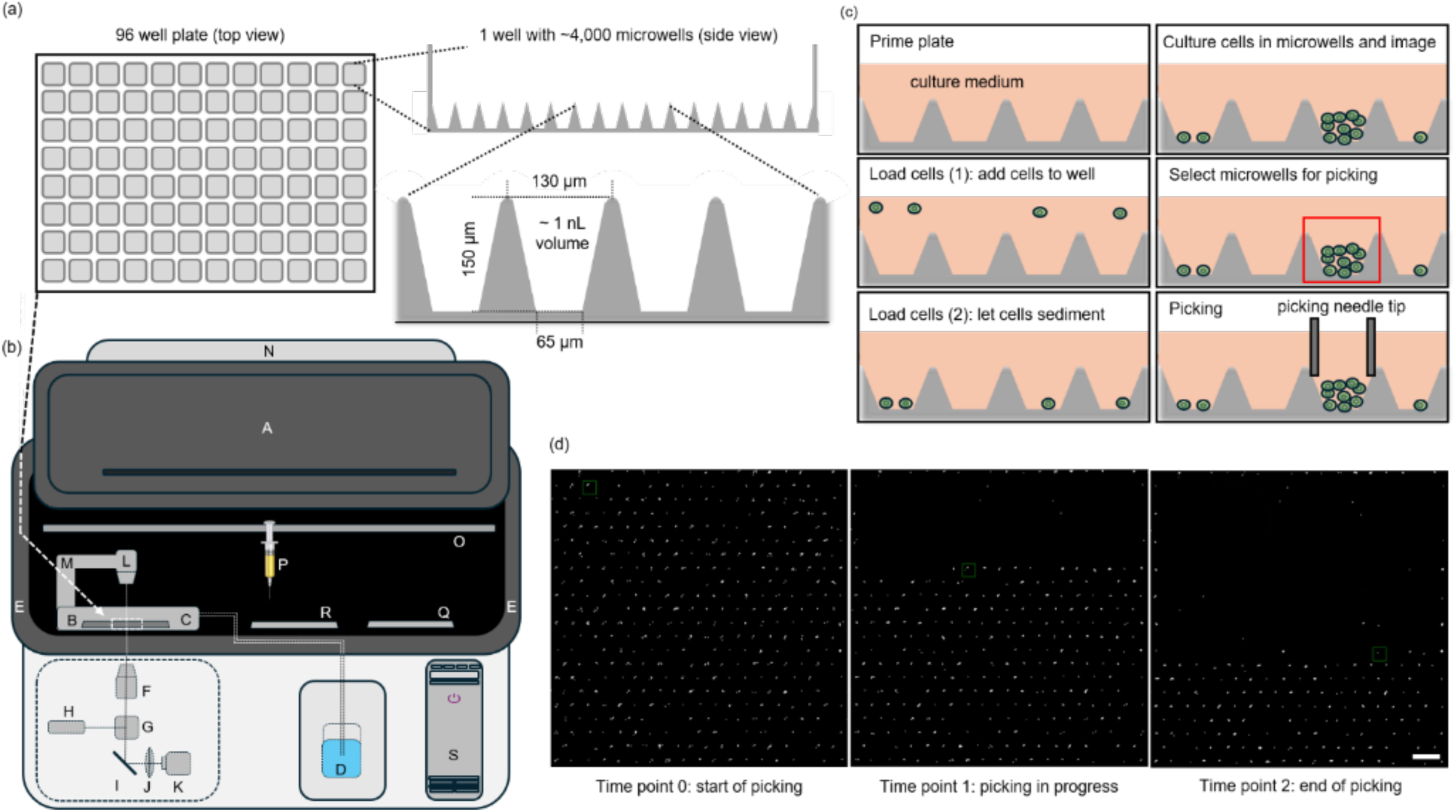
Schematics of microwell arrays and the cloneXplorer. (a) A 96-well plate with an array of conical microwells at the well bottom is shown. (b) The cloneXplorer instrument with its door (A) open is illustrated. A microwell plate (B) is placed inside the stage-top incubator (C) with gas and humidity control (D). The walls of the enclosure (E) are heated to maintain the body temperature. The optical scanner consists of a 4-channel inverted epi-fluorescent microscope (F: 5X objective, G: quad-bandpass dichroic, H: 4-channel LED engine, I: mirror, J: tube lens, K: camera), and transmitted light condenser (L) on a rotational arm (M). During picking, a HEPA filter both maintains a particle-free environment and cools the enclosure. The picker is mounted on a XYZ stage with a syringe (P) effector that is used to transfer liquids between the microwell array (B), temperature-controlled cell deposition plate (Q) and wash plate (R). The instrument is fully automated with software installed on a local workstation (S). (c) Key steps of a workflow are priming an array (filling microwells with culture medium), cell loading, live cell imaging, selection of microwells containing cells of interest and cell retrieval by picking. (d) A picking demonstration to retrieve fluorescent beads from one field of view in sequence is shown at 3 time points. Scale Bar is 200-μm.

### Single cell proliferation

The live cell imager was validated using the NALM6 cell line derived from a patient with acute lymphoblastic leukemia. We optimized the cell loading procedure using a Poisson distribution model, which predicts that single cell occupancy is maximized when the number of loaded cells equals the number of microwells. Under these conditions, loading 4,000 cells into one well yields ∼1,300 microwells containing a single cell (**Fig. S4**). A fast autofocus algorithm was developed for automated time-lapse imaging. The acquired images were subsequently stitched, analyzed, and compiled for hierarchical gating (**Fig. 2a**, see methods for details on data analysis pipeline). Because the picker tip has a finite diameter, it cannot access microwells nearest the well perimeter. Consequently, the working region is limited to the central 75% of the well footprint, which permits ∼2,800 microwells to be accessed by the picker.

**Fig. 2.**
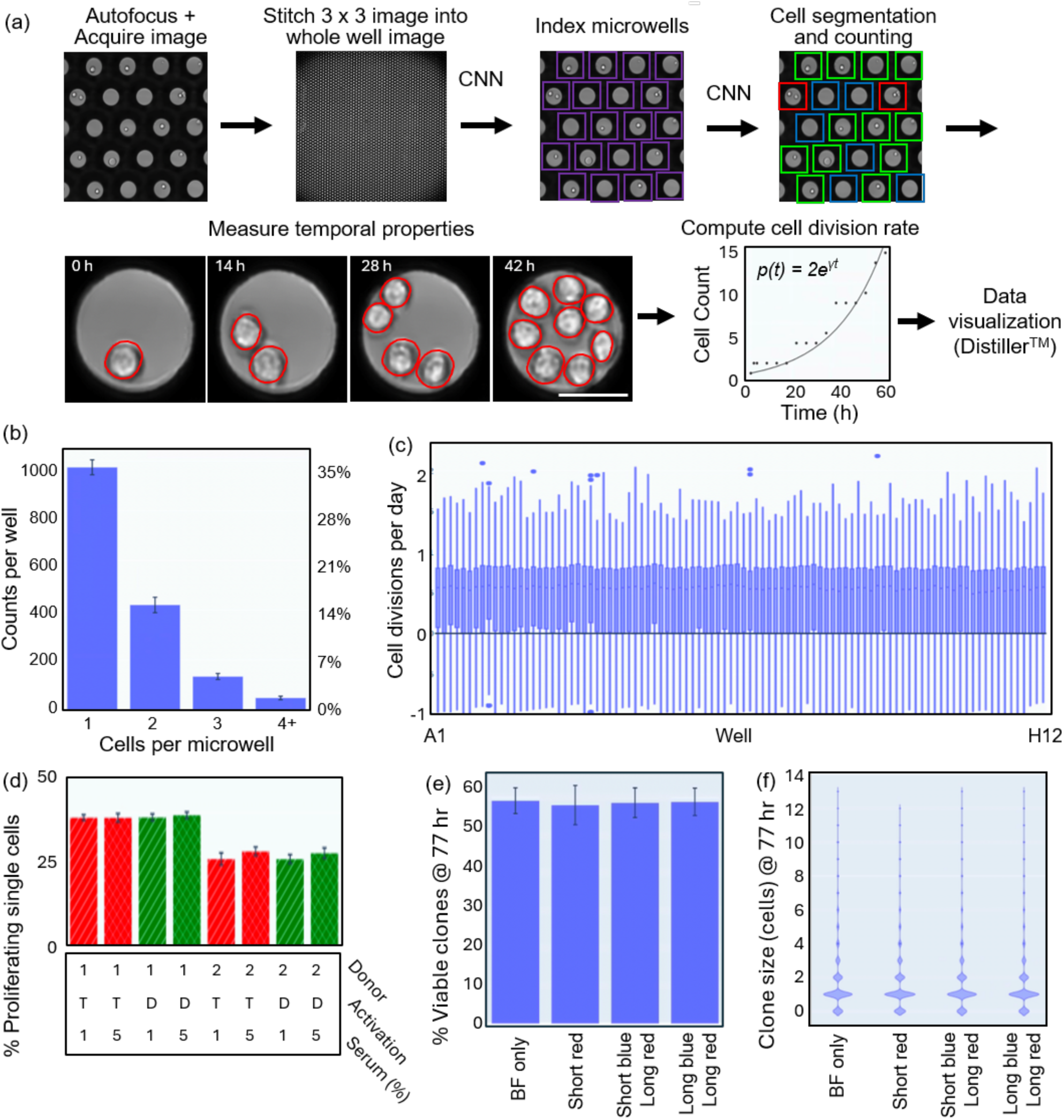
Measurements of single cell proliferation. (a) An overview of the image analysis pipeline is provided. (b) The mean and standard deviation of the number of microwells with 1, 2, 3, and 4+ cells are shown. The percentages are calculated relative to the ∼2,800 microwells imaged per well. (c) Boxplot shows the cell division rates for each well of the plate. In total, the growth measurements for 93,081 microwells containing a single NALM6 cell at the initial timepoint are reported. (d) The fraction of proliferating T cells in two donor samples (Donor 1 and 2) is plotted versus activation matrix (T = Miltenyi Transact CD3/CD28, D = ThermoFisher Dynabeads CD3/CD28) and human AB serum percentage. The mean values for n=4 wells are shown with error bars representing the standard deviation. (e) The mean percentage of viable clones at 80 hours and their standard deviations are shown for donor T cells exposed to different levels of fluorescent light, including 390 nm excitation for 0.05s (short blue) or 0.2s (long blue) exposure, and 565 nm excitation for 0.8s (short red) or 3.2s (long red) exposures. Microwells loaded with a single T cell are shown and viable clones are defined as T cells that divided at least once. (f) Violin plot showing the clone size distribution in terms of the number of cells per microwell for the same conditions as (e). Scale bar is 50 μm.

A user interface (Distiller™) was designed to read the metadata, to graphically visualize all parameters, and to derive (sub)populations by hierarchical gating. Datapoints remain connected to the underlying raw images through time-lapse movies of specific microwells **(Movie S3)**, which are generated on demand to spot check and adjust gate positions. **Fig. 2b** shows a bar chart generated in Distiller, graphing the number of microwells containing 1, 2, 3, or 4+ cells at the initial time point. Loading statistics correspond to the expected Poisson distribution (**Fig. 2b, S4-5**) and is homogeneous across the plate, as indicated by tight error bars. Cell division rates calculated from time-lapse movies of 4-day NALM6 cultures are shown in **Fig. 2c** with a median of around 0.8 cell divisions/day. The wide distribution of cell division rates within each well demonstrates that even established cancer cell lines consist of sub-populations with heterogeneous growth behaviors. Importantly, the growth rate distribution is uniform across the plate even at the edge and corner wells. This phenomenon can also be seen in a map of the cell count for every microwell imaged at 8 hour intervals for 80 hours (**Movie S4**, **Fig. S6**).

To establish suitability of the platform for experiments with primary human cells, we analyzed the proliferation rates of freshly thawed T cells derived from healthy donors stimulated by different anti-CD3/CD28 activation matrices from two vendors and exposed to human antibody serum percentages of 1% vs 5%. **Fig. 2d** indicates that the fraction of proliferative clones depends on the donor but not on the serum percentage or the type of activation matrix (see also **Fig. S7** and **Movies S5-S7**). **Fig. 2e** shows a corollary experiment, in which freshly thawed CD3/CD28 activated T cells were monitored during exposure to different amounts of fluorescent light, and we found no evidence of phototoxicity even after repeated 0.2s exposures to 390 nm light (long blue) and 3.2s exposures of 565 nm light (long red).

Interestingly, the number of proliferating T cells in all samples tested never exceeded 50%, which is in accordance with published proliferation rates of primary human T cells in single cell^10^ and bulk cultures^11^. Final clone sizes after 3 days of culture in microwells starting with single cells is shown in **Fig. 2f**. Importantly, there is no notable phototoxicity, as indicated by the absence of reduction in cell growth due to exposure from fluorescent light.

### Cell line development

To establish suitability of the platform for monoclonal cell line development, we seeded a mixture of mCherry-K562 and NALM6-GFP cells at low density (30,000 cells/mL resulting in 3,000 cells/well) into the microwell array. Cells were continuously imaged at 4 hour intervals for 4 days in brightfield and fluorescent channels. Microwells that contained single cells at the first imaging time point (green rectangle in **Fig. 3a** left panel) were selected with a size gate adjusted to remove debris (small objects classified as cells) and potential artifacts (objects >20 µm). Cell fluorescence was then used to select subpopulations of microwells containing either GFP+/mCherry-cells (solid red rectangle), or GFP-/mCherry+ cells (dashed red rectangle, **Fig. 3a** middle panel). The gates for cell selection were drawn manually to achieve good separation of GFP+ and mCherry+ cells, however these methods can be automated. Lastly, microwells were gated for high growth rates (black rectangle in **Fig. 3a** right panel). As an output of this gating scheme, the Distiller software generated a picking list from which individual microwells were manually selected for isolation and saved in a format that is readable by cloneXplorer.

**Fig. 3.**
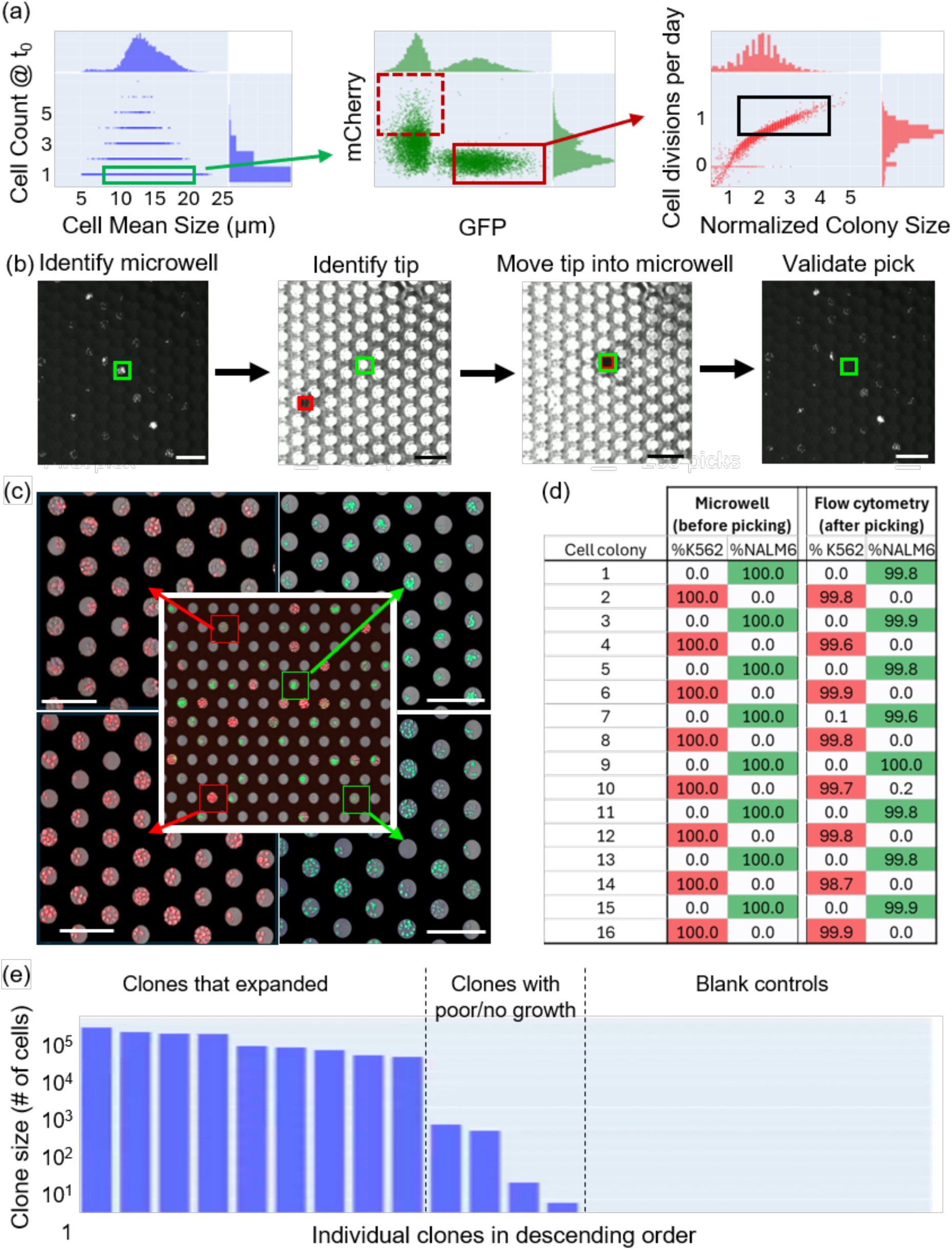
Cell line development workflow. (a) Hierarchical gating scheme to select fast proliferating GFP+ or mCherry+ clones with (b) the main steps of the cell picking process highlighted. (c) A mix of cells with genetically encoded mCherry-K562 (red) or GFP-NALM6 (green) was seeded into a microwell array and allowed to expand (center image). Cells from individual microwells were retrieved for further expansion and seeded into an array (4 corner images). (d) Confirmation of purity of picked clones in a representative sample of 16 out of the 88 colonies isolated. (e) Retrieval and expansion of T cell clones.

**Fig. 3b** illustrates the computer vision pipeline for automatically guiding the picker tip into the microwell of interest (green rectangle). Due to the pronounced optical background of the microwell array in transmitted light images, custom methods were needed to implement computer vision guided motion control. Our optimized method involves gradual lowering the picker tip until it gets near the surface and first becomes visible, allowing it to be identified in XY for the first time, as indicated by the red rectangle in **Fig. 3b**. After initial identification, the tip is moved laterally into the desired microwell using the same models as feedback for position control. The tip height within the selected microwell is then adjusted with another classifier, which establishes a tight seal with the microwell rim. After collecting the content of an individual microwell, the syringe moves to a user-defined well in the deposition plate where the collected cells are deposited. In between each pick, the tip is washed with a combination of ethanol, bleach, water, and cell culture media.

To assess cell cross-contamination risk and the efficiency of clonal outgrowth, we retrieved 88 microwells from the de-mixing experiment. Clones were picked in alternating fashion (GFP+, mCherry+, GFP+, etc.) and grown for 9 additional days until the cell populations exceeded 40,000 cells on average, corresponding to ∼1.2 cell divisions per day through the life of the clone (4 days in microwells + 9 days in the deposition plate). The total number of cells for each clone was measured by flow cytometry and by imaging again in microwells (**Fig. 3c, d**). We found that 84 of 88 clones expanded to a population of greater than 1,000 cells (range 1,400 – 126,000) with no observable contamination detected between subsequent picks, indicating that both the tip washing procedure and the cell identification pipelines perform as expected. The life of each cell from initial seeding through cell retrieval is recorded, with one example of a K562 clone from single cell through multiple days of expansion and retrieval shown in supplementary materials (**Movie S8**).

To test suitability of cloneXplorer for clonal selection of viable primary cells, we loaded freshly thawed human T cells and CD3/CD28 activation beads into a microwell array and imaged at 3-hour intervals for 5 days. Clonally expanded T cells were selected for retrieval while empty microwells randomly distributed in the picking order served as the negative control (blank). Retrieved cells were re-activated with CD3/CD28 beads and cultured for additional 2 weeks followed by cell counting with flow cytometry. **Fig 3e** shows that 9 out of 13 clones expanded while the blank controls were negative, demonstrating usability of the cloneXplorer in cell line development applications with primary cells. An online version of this dataset is hosted at the BioImage Archive^12^.

### Antigen validation workflow

Next, we established a workflow for screening functional TCR-epitope interactions using engineered Jurkat reporter T cells in co-culture with K562 APCs. Jurkat cells were engineered by knocking out the endogenous TCRs, introducing an NFAT-GFP reporter together with CD8 expression cassette, followed by introducing a CMV-specific TCR to monitor immune activation. In parallel, K562 cells were transduced with a pooled library of 100 cancer-associated neoantigen peptides as a trimeric construct peptide-beta-2-microglobulin-HLA-A:02:01 (**Fig. 4a**). When the Jurkat and K562 cells are co-cultured, the NFAT-reporter becomes activated and GFP is expressed only if the TCR binds to the cognate antigen presented by the matching K562 cell present in the same microwell. This Jurkat-K562 artificial system allows a library of antigens to be screened against a single TCR or library of TCR candidates.

**Fig. 4.**
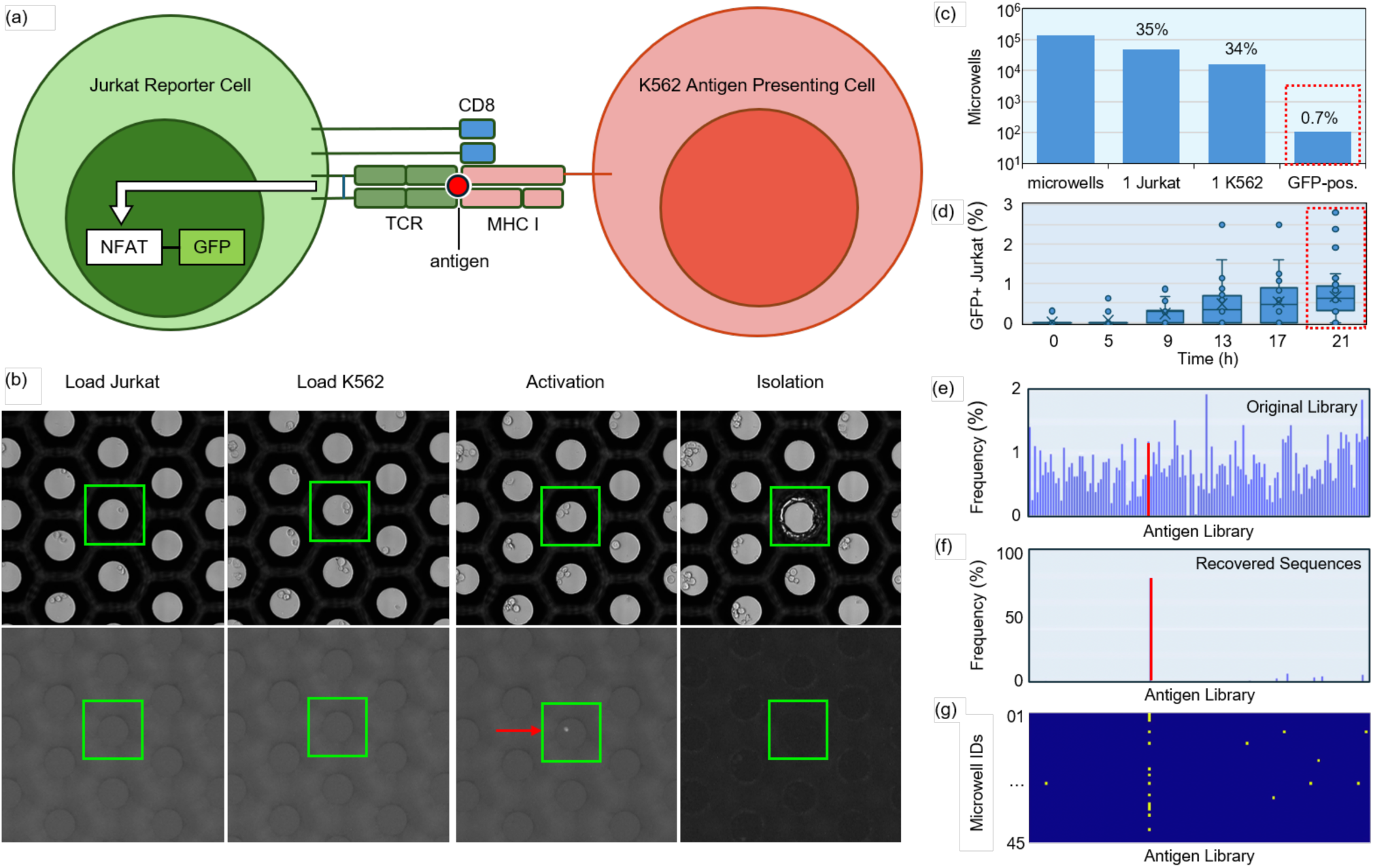
Antigen-specific TCR sequence identification. (a) Illustration of the TCR-antigen cell-based screening platform. Jurkat cells were engineered to express an NFAT-GFP reporter, CD8, and a TCR against a CMV peptide. The white arrow indicates NFAT-reporter activation in the form of a bright GFP+ signal. K562 cells were engineered to express antigens conjugated with beta-2-microglobulin-HLA-A 02:01 complex. (b) Representative images of a microwell at different steps of the workflow. First, Jurkat cells were loaded and imaged. Next, K562 cells were loaded followed by time lapse imaging. Microwells with GFP expression in single Jurkat and K562 cells were identified and selected for retrieval (red arrow). (c) Cells were loaded into 50 wells. Of the 137,758 imaged microwells, 47,597 containing a single Jurkat cell (∼35%). Of those, 16,070 microwells contained a single K562 cell at the second timepoint, corresponding to ∼34% of the prior gate. Of those, 107 microwells contained cells expressing GFP, corresponding to ∼0.7% single cell pairs. (d) Percentage of microwells containing GFP+ Jurkat cells over time. Shown are boxplots with overlaid individual values. (e) The relative abundance of the 100-plex antigen library expressed by K562 cells, determined by bulk sequencing of the original library. (f) The recovered sequences are enriched for the CMV-expressing member of the K562 library. (g) Plot of recovered sequences of antigens (x-axis) vs. microwell IDs (y-axis).

The basic workflow involves first loading Jurkat cells expressing a specific TCR (here, specific to CMV antigen) into the microwells, followed by full plate imaging to identify microwells with single Jurkat cells. Next, a library of K562 cells was loaded into the microwells, after which the plate was imaged again to identify the Jurkat-K562 single cell pairs used for combinatorial screening of singly presented epitopes. The plate was imaged in brightfield and GFP channels at 4-hour intervals for 24 hours (**Fig. 4b**).

Distiller was used to select positive microwells, which are gated based on the presence of a single Jurkat cell at the first timepoint (∼35% of microwells), exactly one K562 and one Jurkat cell at the second timepoint (∼34% of the prior gate), and microwells that were GFP-at the first timepoint and which became GFP+ positive after 9 hours (**Fig. 4c**). In total, this experiment used 50 wells, of which 16,070 microwells were found to contain single Jurkat and K562 cells (1:1 co-culture). Within this subpopulation, we identified 107 GFP+ microwells, corresponding to ∼0.7% of all single cell co-cultures, which is close to the expected frequency of 1% assuming an equally distributed 100-member antigen library. We also quantified non-specific activation by measuring the percentage of GFP+ microwells present within the first few hours after cell loading, which was 0.02% of the microwells with Jurkat-K562 single cell pairs (**Fig. 4d)**. Cells from GFP+ microwells were retrieved, and peptide inserts were amplified and sequenced using primers flanking peptide inserts. We compared the expression of all 100 epitopes in K562 in the bulk pool measured by bulk sequencing (**Fig. 4e**) with the relative frequency of epitopes after enrichment by our functional screen (**Fig. 4f**), where we observed that the CMV-epitope accounted for roughly 80% of the recovered sequences. **Fig. 4g** shows the specific antigens identified in each of 45 analyzed microwells. An online version of this dataset is hosted at the BioImage Archive^12^. Potential screening throughput is discussed in the supplementary materials.

### Cytokine detection

A widely used biomarker of T cell activation is cytokine secretion. Here, we used a fluorescent linked immunosorbent assay, commonly known as FluoroSpot, to detect IFN-γ secretion in co-cultures of single T cells with various types of APCs. The optical sensing range and limits of detection were evaluated by loading IFN-γ capture beads in the microwells, exposing them to different concentrations of recombinant IFN-γ, followed by washing and staining with fluorescent detection antibodies. The sensing range is linear across 3 orders of magnitude with a lower limit of detection of 10 pg/mL (**Fig. S8**). There are several drawbacks of using cytokine capture beads, such as need to load beads at relatively high concentration to ensure that >95% of microwells have at least one bead (**Fig. S4**). Also, some cells, such as monocyte derived dendritic cells (DCs) used as APCs, phagocytose the beads. To avoid these issues, we developed a method to immobilize capture antibodies on the microwell walls and monitor IFN-γ accumulation during the experiment (**Fig. 5a-b)**. Notably this method does not require any rinsing steps, since the detection antibodies are always present in the culture media and continually accumulate on the microwell walls over time.

**Fig. 5.**
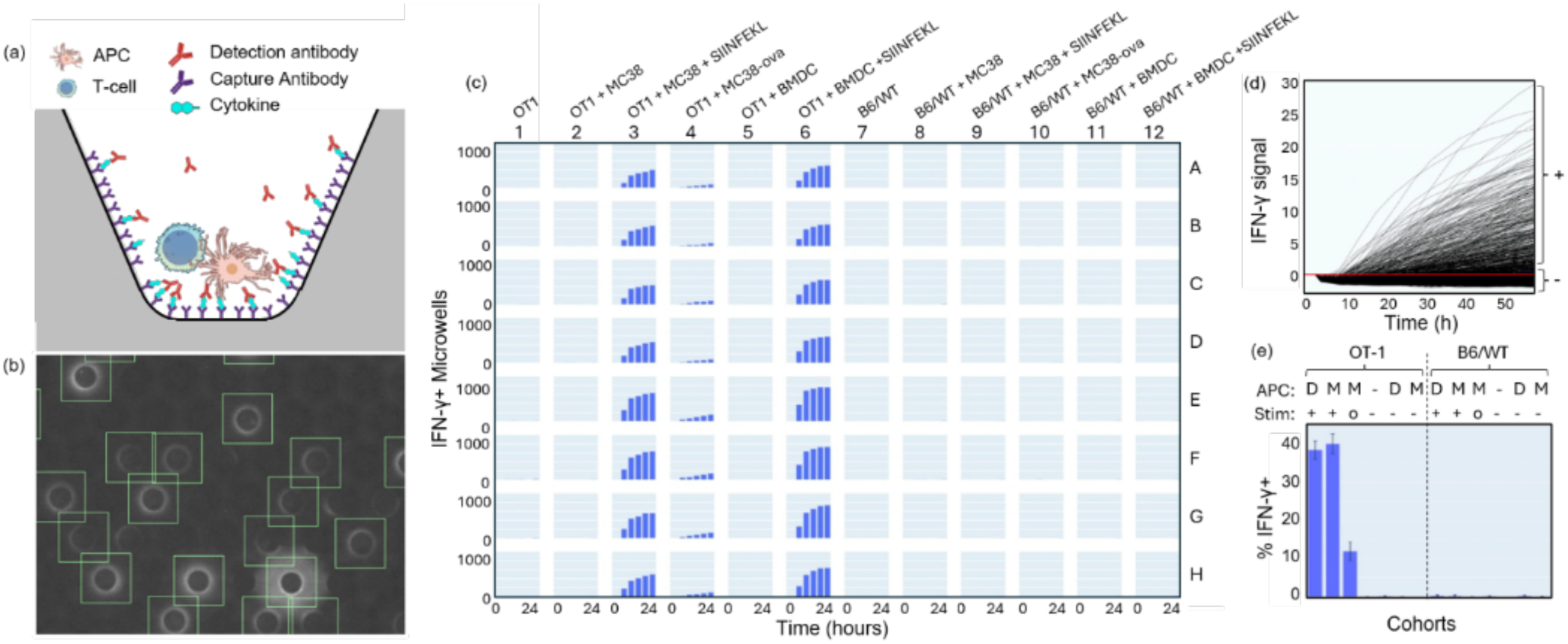
Cytokine detection platform. (a) Cytokine detection by FluoroSpot. (b) Fluorescence image showing characteristic secretion rings of microwells containing IFN-γ secreting cells. The green boxes depict microwells with accumulated IFN-γ. (c) Bars correspond to the number of microwells with accumulated IFN-γ in each well of the plate over 24 hours. (d) The change in the intensity of accumulated IFN-γ over 60 hour experiment in one well of the plate is plotted for all 1,052 microwells containing single T cells. (e) The percentage of microwells with accumulated IFN-γ is shown for 12 experimental conditions. The APC legend defines D as BMDC, M as MC38 cells, and “–” means that APCs were absent. The stimulation legend (Stim) defines “+” as soluble SIINFEKL-peptide, “–” as the absence of soluble peptide, and “o” indicates the use of MC38-ova to stimulate the T cells. The effectors were T cells derived from either OT-1 or B6/WT mice.

To benchmark the platform against a well-established *in vivo* model, we compared ovalbumin antigen specific CD8+ T cells from OT-1 mice^13^ with T cells from B6/WT mice. Both groups of T cells were seeded in co-cultures with APCs, which included either bone marrow derived DCs (BMDC) loaded with the antigenic ovalbumin peptide SIINFEKL or MC38 colon cancer cells overexpressing ovalbumin (MC38-ova). Negative controls for the APCs were BMDC or MC38 cells without peptide loading. Of note, adherent MC38 cancer cells grow as spheroids in our microwells, since the microwell walls are coated with anti-adherence solution (**Movie S9**). The number of IFN-γ+ microwells in each well is plotted for the first 24 hours (**Fig. 5c**, **S10**), with **Fig. 5d** indicating that cytokines secreted by OT-1 cells can be detected in the first 8 hours, and the level of secretion continues to increase for at least 2 days (**Movie S10**). These experiments measured secretion continuously for 2 days, however this assay can be extended to quantify slower activation dynamics, such as when naïve T cells are first exposed to an antigen matching their TCR. The IFN-γ signal at each timepoint was assessed by averaging the fluorescent intensity within an annulus that encloses the walls of the microwells (secretion ring). A secretion map of every microwell over time is shown in supplementary materials (**Movie S11**). **Fig. 5e** shows that, in the absence of stimulation, around 1 microwell per well has detectable accumulation of IFN-γ (0.03% of microwells). Hundreds of positive microwells were detected in each well containing stimulated T cells, corresponding to ∼40% of the microwells with co-cultures were activated in the presence of soluble peptide and ∼10% of microwells with co-cultures activated with over-expressed cell-based antigen presentation. An online version of this dataset is hosted at the BioImage Archive^12^.

### Monoclonal antigen-reactive T cell line development and sequence identification

A workflow was developed to identify and retrieve antigen-reactive T cells from Peripheral Blood Mononuclear Cell (PBMC) samples (**Fig. 6a**). In a bulk enrichment step, we add synthetic Melanoma Antigen Recognized by T cells (MART1) peptide to PBMC cultures leading to presentation of the peptide by APCs and then expansion of antigen-reactive T cells. For the melanoma antigen MART1, we typically achieve 100-fold enrichment using PBMCs from healthy donors after 7 days of culture (**Fig. 6b-c**). In parallel with the enrichment step, we generate DCs from monocytes of the same donor, and pulse them with peptide before combining them in microwells with the enriched T cells to select clones for isolation by measuring IFN-γ secretion. Although there is background secretion in some of the control groups (e.g. green bars in **Fig. 6d**: T cells + unpulsed DC), the number of IFN-γ+ microwells is significantly higher in the experimental group (red bars: T cells + DC + MART1-peptide). Anti-CD3/CD28 antibody complexes used as the positive control confirm that a high number of T cells produce IFN-γ in response to non-specific activation (blue bars, **Fig. 6d**). After 5-6 days culture in microwells, we stain the cells with anti-CD8 and anti-CD4 antibodies making use of our deep microwells to enable washing steps without disturbing the cells trapped at the bottom. We selected clones (**Fig. 6e**) based on detectable IFN-γ secretion, T cell proliferation, and CD8 expression. Our detection method avoids issues with heterogeneous background fluorescence by detecting the characteristic “ring-shaped” secretion pattern on the walls of the microwells, rather than relying on absolute fluorescent intensity to indicate positive events (Fig. 6e, left). **Fig. 6f** shows representative images of a microwell seeded with 1 T cell and 2 DCs. This T cell starts secreting IFN-γ at ∼24 hours and expands into a CD8+/CD4-colony (left panel). The right panel in **Fig. 6f** depicts images before and after picking, revealing that the selected CD8+ colony was removed by the picker. In total, 60 colonies of T cells were retrieved, clonally expanded, and quantified by flow cytometry with fluorescent MART1 tetramer stain. One example of a MART1-positive T cell clone is depicted in **Fig. 6g**.

**Figure 6.**
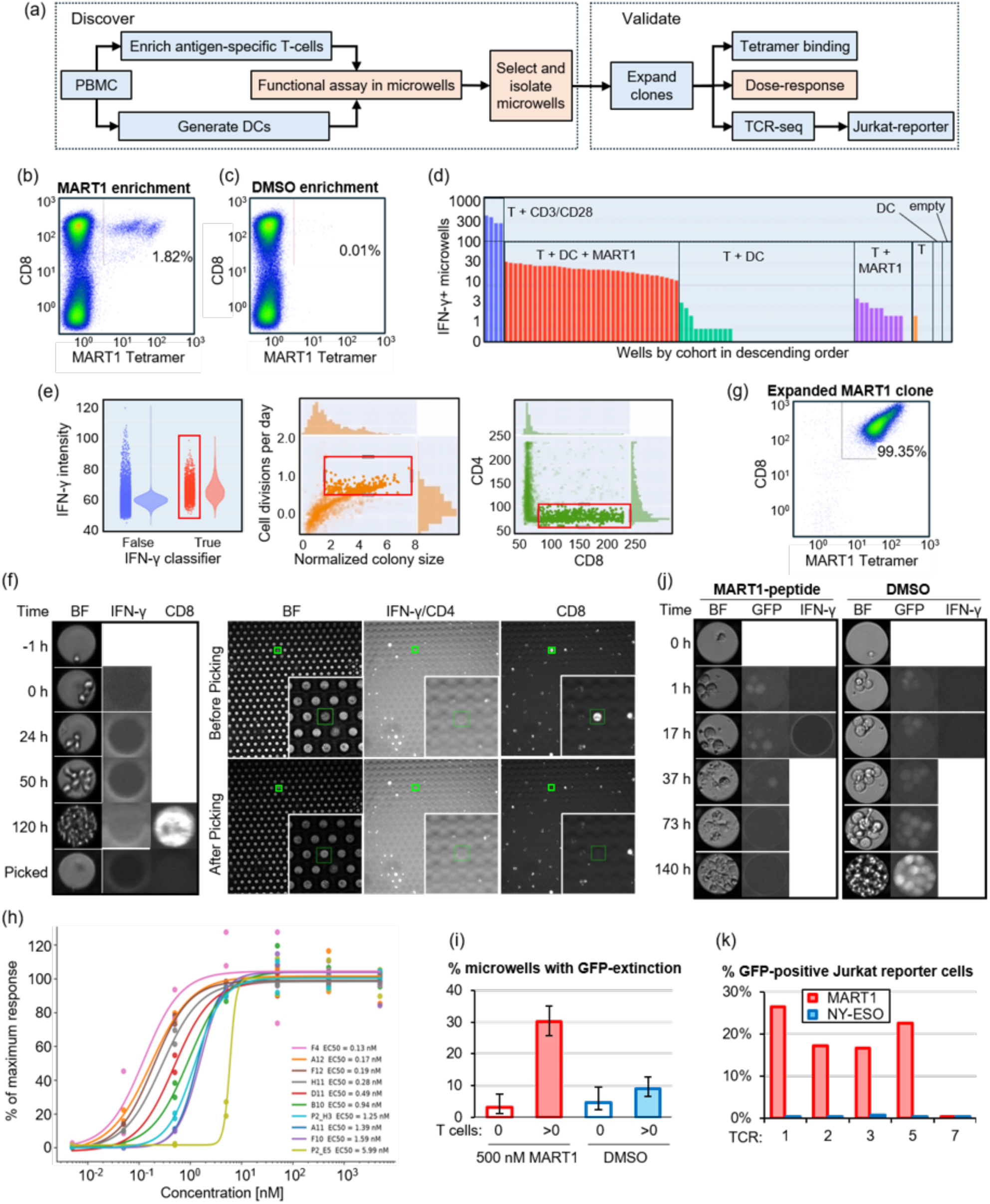
Generation and validation of antigen-reactive T cell lines. (a) The overall workflow is illustrated. (b-c) Flow cytometric plots of T cells stained with anti-CD8 antibody and MART-1 tetramer after enrichment with either MART-1 peptide (b) or DMSO (c). (d) Number of IFN-γ+ microwells per well, with color denoting the condition: T cells with anti-CD3/CD28 antibodies (blue, positive control), T cells with dendritic cells and MART-1 peptide (red), T cells with unpulsed dendritic cells (green), T cells with MART-1 peptide (purple), T cells (orange), dendritic cells or no cells. (e) Hierarchical gating strategy to identify microwells with an IFN-γ signal (left), containing proliferating cells (center) and containing CD8-positive cells (right). IFN-γ is detected using a classifier that recognizes the characteristic pattern of secretion rings (x-axis) and is associated with higher fluorescence intensity measurements in the secretion ring area (y-axis). (f) Example of one clone pick. Left panel: Images of a selected microwell at multiple time points in brightfield (BF) and fluorescence (IFN-γ and CD8). BF is used to assess initial conditions (-1 h: 1 T cell loaded, 0 h: additionally 2 DCs loaded) as well as growth rate. Right panel: Images of the array before picking and after picking are shown in all channels. The disappearance of cells after picking is noticeable in the CD8 channel. (g) Flow cytometry plot of the cells picked in (f) that were further expanded before analysis. (h) Antigen-reactivity validation of 10 expanded cell lines by re-challenging them with MART1-peptide pulsed APCs in a dose-response experiment. IFN-γ secretion normalized to maximum secretion of each cell line is shown across different peptide concentrations, along with 4 parameter logistic regression analysis to determine EC50-values. (i) MART1-dependent killing of peptide pulsed APCs by cell line F12. 500 microwells were analyzed for each MART1 or DMSO-pulsed APCs and separated by the presence (>0) or absence (0) of T cells. Error bars indicate 95% confidence intervals. (j) Representative microwells loaded with 1 T cell of the F12 cell line (0 h) and 3 GFP-expressing APCs (1 h) that were pulsed with 500 nM MART1-peptide (left) or DMSO (right). MART1-pulsed APCs are killed and activate the T cell to secrete IFN-γ and proliferate (left), whereas DMSO-treated APCs do not activate the T cell and expand. (k) GFP expression of Jurkat NFAT-reporter cell lines expressing TCRs 1, 2, 3, 5 or 7, and co-cultured with either MART1-peptide or NY-ESO-peptide expressing APC.

To confirm reactivity towards MART1, 10 expanded cell lines were re-challenged in an IFN-γ secretion assay with HLA expressing K562 APCs pulsed with MART1-peptide titrated from 5,000 – 0.005 nM. The number of IFN-γ+ microwells for each T cell line was MART1 concentration-dependent with EC50-values ranging from 0.1 to 6.0 nM (Fig. 6h). We further characterized three cell lines for cytotoxicity by measuring the disappearance of GFP-labeled target cells in microwell co-cultures. Cell line F12 is shown as example in Fig. 6i, in which 30% of T cells were observed to kill one or more GFP-labeled K562 APCs. Representative microwells of a T cell killing 3 GFP-labeled MART1-pulsed APCs is depicted in Fig. 6j (Movie S12) as a comparison against DMSO-pulsed APCs that did not solicit a cytotoxic response, as depicted in Fig. 6j (Movie S13). After functionally validating these T cell lines and characterizing their antigen reactivity, we sequenced their TCR-alpha (TRA) and beta (TRB) chains, leading to the discovery of 4 unique TCRs, and 3 shared by 2 cell lines (translated amino acid sequences of CDR3 regions are shown in Table 3). A comparison of the EC50-values for each cell line reveals that cell lines sharing a TCR differed in EC50 by only 1.1- to 2.2-fold, compared with a median 4.3-fold difference between cell lines expressing different TCRs (exact permutation test, p = 0.009; rank-based sensitivity analysis, p = 0.068). Given that only three TCRs were represented by more than one cell line, this indicates an association between TCR identity and functional avidity. Five identified TCRs were expressed in Jurkat NFAT-GFP reporter cells and cocultured with APCs expressing either MART1 or the unrelated NY-ESO single chain trimer (SCT). TCRs 1, 2, 3, and 5 were strongly activated by MART1, but not by NY-ESO (Fig. 6k), whereas TCR 7 was not reactive. We noticed that cell line P2_H3 from which TCR 7 was recovered expressed two TRA mRNAs and hypothesize that the non-productive TRA was chosen for expression studies (Table 3). Taken together, these data demonstrate that we can identify, isolate, and expand antigen-reactive T cells that are present in PBMCs at frequencies as low as 0.01%. An online version of this dataset is hosted at the BioImage Archive^12^.

**Table 1.** Comparison of commercial systems capable of analyzing cell function at multiple time points with ability to retrieve clones of interest for applications in cell line development and immune discovery.

|  | **Bruker** | **Cell Microsystems** | **Sartorius** | **Celldom** |
|---|---|---|---|---|
| Instrument | Beacon | CellRaft Air | CellCelector | cloneXplorer |
| Consumable | OptoSelect 3500 | CellRaft | Flex S200 | cloneXpress |
| Number of samples | 4 | 24 | 6 or 24 | 96 |
| Partitions per experiment | 14,000 | 154,000 | 100,000 | 380,000 |
| Single cell occupancy | > 80% | < 40% | < 40% | < 40% |
| Partition aspect ratio (H/W) | > 10 | 0.5 | 0.5 | > 2 |
| Loading Time | 1 hr | 10 min | 10 min | 10 min |
| Environment control | Yes | No | No | Yes |
| Media exchange | Yes | No | No | Yes |
| Small molecule drug screens | Yes | No | Yes | Yes |
| Reference | Ref. <sup>24</sup> | Ref. <sup>14</sup> | Ref. <sup>15</sup> | **This work** |

**Table 2.** Datasheet for the cloneXplorer.

|  | **Specifications** |
|---|---|
| Consumable | 96 well plate (SLAS) |
| Imaging Fields (9 per well) | 864 |
| Image pixel size | 0.5 µm |
| Full plate scan BF only | 15 min |
| Full plate scan BF + 4-FL channels | 30 min |
| Autofocusing | every imaging field |
| Temperature Control | +/-0.2<sup>o</sup>C |
| Pickable microwells per well | 2,800 |
| Total microwells per well | 4,000 |
| Humidity Control | >90% R.H. |
| CO<sub>2</sub> control | 0 – 10% |
| O<sub>2</sub> control | 0 – 20% |
| Full plate image analysis | 60 min |
| Picking time without rinsing | 30 seconds |
| Picking time with rinsing | 2 - 5 minutes |
| Picking success rate | >90% |

**Table 3.** Characterization of 7 TCRs found in 10 cell lines. EC50 values from 4 parameter logistic regression (Fig.6h). Amino acid sequences of CDR3 regions for TRA and TRB. Cell line P2_H3 had transcripts of 2 TRA chains; the underlined sequence was tested in Jurkat cells and found not to recognize MART1 when combined with the listed TRB (Fig.6k). n.a. not analyzed.

| TCR | cell line | EC50 | CDR3 (TRA) | CDR3 (TRB) | Jurkat-reporter |
|---|---|---|---|---|---|
| 1 | F10 | 1.59 nM | CAVRNNDMRF | CASSQDAIGQPQHF | reactive |
|  | A11 | 1.39 nM |  |  |  |
| 2 | F12 | 0.19 nM | CVVNMWSSNTGKLIF | CASSFGNEQFF | reactive |
|  | A12 | 0.17 nM |  |  |  |
| 3 | D11 | 0.49 nM | CAVNSGGGADGLTF | CAIRRLAGESNGELFF | reactive |
| 4 | H11 | 0.28 nM | CAPNSGGGADGLTF | CASSFQGSGTEAFF | n.a. |
|  | F4 | 0.13 nM |  |  |  |
| 5 | B10 | 0.94 nM | CALEAGTALIF | CASRPPGLAGDLSYNEQFF | reactive |
| 6 | P2_E5 | 5.99 nM | CAADPGFQKLVF | CASSLGKAADTQYF | n.a. |
| 7 | P2_H3 | 1.25 nM | CAEIPINQGGKLIF,<br>CAVTGDYKLSF | CASRTGTGAGEQYF | not reactive |

## DISCUSSION

The ability to track individual cells over days and isolate desired cells for monoclonal expansion solves a major pain point for target validation workflows that require antigen-reactive T cell populations for live cell functional assays. In a typical approach, T cells sourced from peripheral blood are isolated either by labeling and sorting of cells with peptide-HLA tetramer complex or by exposing cells to the antigen and sorting them based on activation markers, such as CD69, CD137, or IFN-γ secretion. The isolated cells are then analyzed by scTCR-seq to quantitate the alpha and beta chains, generating a list of candidate TCRs^1^. Individual TCRs are then cloned into reporter cell lines for use in validation studies. The drawback of this approach is that the viable, stimulated T cells are being destroyed during scTCR-seq analysis, which could be avoided if it were possible to efficiently discover and clonally expand the desired T cells in a single, unified workflow.

With this goal in mind, we set out to develop a platform with sufficient scale to identify and isolate antigen-reactive T cells from peripheral blood, which can be rarer than one in a million. We chose microwell arrays^14–18^ as opposed to droplet encapsulation^19–23^ or other microfluidic devices^24,25^, because we believe 96-well plates are the most suitable platform for live cell assays, and they are easiest to automate with standard liquid handlers, plate grippers, and incubators. We favor stochastic cell loading methods^3,15–18,20,21,23,26,27^ over deterministic cell manipulation techniques^6,22,24,25,28^ for reasons of simplicity, biocompatibility, and parallelizability (**Table S1**). Although we cannot deterministically control cell loading, we achieve a high number of experimental conditions with single cells (e.g. ∼1,000 microwells with single cells per well), which is enabled by the high density microwell arrays with empty and overloaded microwells used as the built-in controls. This platform is designed to flexibly interrogate approximately 100,000 single T cells in an experiment and collect time-resolved, multiparameter phenotypic data from each microwell for more than a week. The 96-well format allows testing of up to 96 conditions (e.g. candidate antigens or drugs) in parallel. Additionally, antigens can be pooled in a primary screen (realistically up to ∼10/well) and then de-multiplexed in a secondary assay, increasing throughput to ∼1,000 epitopes. We optimized imaging speed to enable full plate scans in 4-channels in less than one hour (**Table S2**), which can further increase throughput when used in combination with a robotic plate handler.

To improve the cell picking process, we designed conical microwells that perfectly match the diameter of the syringe tip, serving as a self-guiding tip positioning system that protects cells by both guiding the tip into the target microwell and preventing the tip from touching the cells at the bottom. We developed picking protocols that include tip sterilization and washing procedures in between picks, enabling automated isolation of cells from ∼192 individual microwells in 8 hours. High aspect ratio conical microwells also yield major workflow improvements by enabling the exchange of solutions during stimulation and immunostaining without disturbing the cell at the bottom. Reduced convection in microwells also improves signal to noise ratio in cytokine secretion assays. Finally, the ability of deep microwells to hold hundreds of cells increases the probability that cells will expand after isolation.

The cloneXplorer platform is ideally suited for analyzing suspension cultures, spheroids, and organoids; however, conical microwell arrays are less optimal for analyzing adherent cells, which can migrate along the walls and into adjacent microwells. To better confine cells, it is possible to differentially pattern cell attachment proteins at the microwell bottoms and backfill the walls with non-adherence coating; however, some applications are inherently better suited to microfluidic perfusion platforms. Additionally, while our open well platform enables easy access to the cells, it is unable to perform chemical perturbations with precisely controlled timing like perfusion-based systems. For these reasons, we have focused our efforts on immunological screens and in studying cell-cell interactions between suspension cultures.

One of the challenges of microwell arrays is fully utilizing the enormous datasets containing time-correlated morphological and fluorescent measurements. Next generation sequencing technologies faced a similar data challenge in the early days, and it benefited from efforts by the broader scientific community to make standardized datasets available to the public. Towards that end, we have posted raw image datasets and related metadata for the experiments of **Figs. 2**, **4**, **5**, and **6** of the main text in BioImage Archive^12^.

Based on our experiments with clone isolation and sequencing from the CMV-TCR Jurkat reporter cell line, we have shown that the cloneXplorer platform can identify matched TCR-antigen pairs from high complexity libraries (up to 100,000). The simplicity, throughput, and the possibility to enable library-on-library screens (e.g. 100 TCRs against 1,000 antigens) compare favorably to published methods for antigen epitope screening^19,20,27,29–31^.

This work presents significant advances in integrating multiple time-resolved functional measurements (growth, secretion, surface markers) into robust workflows that combine the identification, isolation, and expansion of rare, antigen-reactive T cells derived from peripheral blood. We are currently using this workflow for discovering and validating TCRs reactive to cancer-specific neoantigens presented in MHC class I complexes for use in personalized cancer therapy; however, we have only scratched the surface of potential applications, ranging from cell line development, antibody discovery, low input cell sorting, isolation of rare drug-resistant cancer cells, and as a tool for studying epigenetic drift in cell populations over generations. We are currently working on several of these applications. For example, we have successfully used this platform as a low input cell sorter, in which we isolated single cells from cerebrospinal fluid biopsies having less than 1,000 cells in a 3 mL aliquot. The lack of dead volume allows us to identify nearly every cell deposited in the well and isolate cells with defined morphological features. Another example is the investigation of drug resistance in cancer cell lines and primary cancer samples, in which we identified rare cancer cells that survive drug treatment for whole genomics analysis and/or clonal expansion. With future development, we hope that this platform will enable applications in personalized drug screening and even personalized biological medicines.

## MATERIALS AND METHODS

### Consumable

Conically shaped microwell arrays were designed with a 65 μm diameter bottom, center-to-center distance of 130 μm pitch, and a minimum height of ∼150 μm (**Fig. 1a, S1-2**). The master mold to produce these conical shapes was developed using a proprietary lithography system. The mold was pressed into a plastic bilayer consisting of a deformable thermoplastic film bonded to a rigid 188 μm thick cyclic olefin polymer (COP) film, which is used both as a backstop and as a substrate for high quality optical imaging. The substrate is cut to shape and fastened to an open bottom 96-well plate frame, packaged, and sterilized.

### Instrument

We built a time-lapse imaging platform with a clone picker and laminar flow hood to maintain sterility during picking (cloneXplorer, **Fig. 1b, S3**). The imaging module consists of an inverted microscope with transmitted light (730 nm) optical condenser for brightfield imaging, wide-field epifluorescence (390 nm, 488 nm, 565 nm, 630 nm excitations), 5X microscope objective, 4 channel LED engine, quadruple bandpass dichroic cube, and 20 MP camera. An on-stage incubator was used to control CO_2_, O_2_, and humidity. The walls of the instrument enclosure are heated to maintain the system temperature at 36.5°C ± 0.2°C. The stage-top incubator is mounted on a microscope XY-stage and a Z-stage was used to move the microscope objective for autofocusing. A microinjection syringe mounted on a separate XYZ stage is used for retrieving the contents of specific microwells and transferring them to a deposition plate. To illuminate the sample during the cell retrieval process, a miniature brightfield condenser was built around the syringe tip. The imaging condenser was mounted on a rotational stage, so that it can be moved out of the way during image-guided cell picking. The deposition plate can be cooled down to 0°C to control evaporation during clone picking, and a wash plate is used to sterilize the syringe tip in between picks. Sterility is maintained during the cell picking process by ventilation through a HEPA filter mounted at the ceiling of the instrument.

### Cell Culture

NALM6-GFP cells (purchased from Imanis Life Sciences) were cultured in RPMI (Corning) supplemented with 10% FBS (Corning). Primary T cells and PBMC were purchased from Stemcell Technologies and cultured in RPMI supplemented with 5% human AB serum (Gemini Bio), and 5 ng/mL recombinant human IL-7 and IL-15 (R&D Systems). T cell activation was performed according to the manufacturers’ instructions (Miltenyi Biotech: human T Cell Transact, ThermoFisher: Dynabeads human T-activator CD3/CD28). Lyophilized MART1 peptide (Anaspec) was dissolved in DMSO at 5 mM. DCs were derived from monocytes, isolated from PBMC using EasySep monocyte isolation kit (Stemcell Technologies), and differentiated and matured in AIM V culture medium (ThermoFisher) following published methods^32^. Recombinant human GM-CSF, IL-4, IL-1β, TNF-α, IL-6 and prostaglandin E2 were purchased from R&D Systems.

### Celldom Plate Priming Procedure

The priming procedure employs centrifugation to replace the air inside the microwells with the working fluid of either an anti-adherence rinsing solution (Stemcell Technologies) or a solution containing capture antibodies to functionalize the surface for cytokine secretion assays (see Cytokine Analysis below). All wells are filled with 150-200 µL liquid and then centrifuged at 2,500 g for 40 min with temperature set to 4°C. After centrifugation, the success of the priming procedure can be verified by the lack of microwells with bubbles via microscopy. Primed plates can be stored at 4°C for at least a week.

### Cell Loading

Cells are loaded by dispensing 100 μL aliquots of cell suspension dropwise at ∼5 mm above the liquid surface. A volume of 100 μL/well is used for cell loading with a concentration of 40,000-200,000 cells/mL depending on the desired cell density (see **Fig. S4-5**). After cell loading, the plate is rested for 10 min on a flat surface to allow the cells to sediment into the microwells. The plate is then centrifuged at 300 g for 2 min to transfer any cells trapped on the conical walls to the bottom of the microwells. If loading cells in wells containing 200 μL liquid, the final volume after loading is 300 μL/well. If additional cell type(s) are loaded to establish co-cultures, 100 μL supernatant is removed from each well after each loading step to reset the volume to 200 μL/well.

### Image Analysis (Fig. 2a)

Each well of the 96 well plate is captured as a 3 x 3 image stitch and saved in Open Microscopy Environment Next Generation File Format^33^ (OME-NGFF) using the zarr data structure developed for storing large N-dimensional image data. The data is stored in TCZYX format, where T is the number of timepoints, C the number of imaging channels, Z the number of z stacks, with Y and X representing the dimensions of the stitch. The image analysis pipeline identifies and crops microwells from the stitch, assigns a unique id to each microwell, and records its coordinates at the first timepoint. For subsequent timepoints, the shifts in the cropping parameters are adjusted to ensure that cell statistics and movies are correctly registered in time. Various properties of the cropped microwells are analyzed at each timepoint, including the mean intensity of the crop in all channels (used to eliminate junk and autofluorescence), fluorescent distribution on the walls of the microwells (used for cytokine detection), segmentation masks of the cells in each microwell, and the fluorescent intensity measurements within each mask used for surface marker detection and cell counting. Images are analyzed in real time, so that the data can be used for clone selection at the desired timepoint. All computer vision models were developed and trained in house on internally generated datasets, as previously described^34^. Faster-RCNN models for detecting microwells and its ring-shaped secretion patterns, as well as tip finding algorithms, were developed in pytorch using a ResNet-50 backbone and feature pyramid network. Instance segmentation models for cell and bead masks were developed from Mask-RCNN models using the same model architecture with different head layers for detection vs. segmentation. Training sets were developed by combining manual and semi-automated annotation strategies. Specifically, a coarse model was first developed by fine tuning an open-source model on ∼1,000 manually annotated images from our experiments. The resulting model was then used to automatically generate a larger dataset of annotations, which were corrected by hand. Using this approach, we built a large, human-supervised, annotated dataset of cell instances that can accurately identify cells in brightfield images. The mask from the brightfield image is used as the template to quantify the mean fluorescent of cells in different channels, allowing us to assign cell types based on surface markers, such as CD4 and CD8.

### Growth Rate Detection

We developed two methods to quantify proliferating colonies. In one approach, we fit the cell trajectory to an exponential growth curve (**Fig. 2a, Movie S3**), where the exponent represents the growth rate in units of cell divisions per day. Positive growth rates indicate growing populations while negative growth rates indicate cell death. This approach accurately identifies the proliferating cell populations in most microwells; however, it was susceptible to imaging artifacts from overlapping cells, apoptotic cells, and other debris, and it did not accurately characterize cell populations that follow abnormal trajectories. To establish a second measure of cell proliferation, we also quantified the mean cell count in the microwell over time divided by the number of cells in the microwell at the initial timepoint. In essence, this is normalized weighted average of the cell count per microwell with values greater than 1 indicating cell growth and values less than 1 indicating cell death. While this parameter was less sensitive to an erroneous cell count at any individual timepoint, it was also less sensitive to hypergrowth owing to the influence of the early time points. By gating based on both parameters, we consistently identified the microwells with highly proliferative cells.

### Cell Picking

A picking list was created in Distiller by filtering populations based on the parameters of interest, and then manually curating the final pick list after reviewing movies from the subpopulations. Upon starting the picking procedure on the cloneXplorer, the imager first moves to the desired microwell, aligns the current image against the previously acquired frame, and identifies the closest microwell based on stored microwell coordinates. Next, the brightfield condenser rotates out of the way and the picker moves into the optical axis and identifies the same microwell for retrieval. The picker consists of a 100 μL volume syringe (WPI) with a replaceable 36G stainless steel needle (WPI) having an outer diameter of ∼110 μm, which is smaller than the microwell opening. Both the tip and the microwell array are flexible, allowing the tip to be guided into the closest microwell of interest. It is only possible to identify the tip when it is near the substrate, which appears as a shaded patch against a high contrast periodic background of microwells. Once the tip is identified in XY, it is moved into the microwell of interest before activating the syringe pump. A typical volume of 1.5 μL efficiently removes the entire contents of the microwell. In between each pick, the tip was washed via a series of fluid exchanges, often starting with ethanol or 10% bleach, and then moving to water, PBS, and then the working fluid for the next pick, which is usually cell culture media.

### Expansion of primary antigen-reactive T cells after picking

Isolated T cells were deposited in U-bottom 96 wells that were pre-seeded with 200,000 irradiated PBMC (iQ Biosciences) in RPMI medium supplemented with 5% human AB serum, 5 ng/mL of each IL-7 and IL-15, 100 U/mL IL-2 (R&D Systems), and 30 ng/mL anti-CD3 antibody (clone OKT3, BioLegend). Cells were fed every 3-4 days with the same culture medium (without anti-CD3 antibody) and passaged into larger wells after significant cell growth was observed.

### Flow Cytometry

For verification of GFP or mCherry expression, K562 and NALM6 cells were labeled with viability dye Viobility 405/520 (Miltenyi Biotech) according to the manufacturer’s instructions. For investigation of antigen reactive T cells, 100,000 – 1,000,000 cells were labeled in 50 µL staining buffer consisting of dPBS (no calcium, no magnesium, ThermoFisher), 0.5% bovine serum albumin (Sigma-Aldrich), 1 mM EDTA (ThermoFisher). Cells were first incubated with blocking reagent Human TruStain FcX (Biolegend) and PE-labeled tetramer (MBL Life science) for 20 min, and then with Alexa Fluor 647-labeled anti-CD8 antibody (BioLegend) and viability dye (Miltenyi Biotech). After washing with staining buffer, cells were resuspended in 100 µL staining buffer and analyzed on a MACSQuant 16 flow cytometer (Miltenyi Biotech).

### Peptide-dose response and EC50 calculation

HLA-A*02:01 expressing K562 Antigen Presenting Cells (Cellecta) were pulsed with 1:10 dilutions of MART1-peptide starting with 500 nM for 2 hours before combining with expanded T cell lines in Celldom plates for IFN-γ secretion analysis.

The number of IFN-γ positive microwells were recorded for individual wells at each of seven antigen concentrations with two replicate wells per concentration. Dose-response relationships were fitted with a four-parameter logistic model parameterized on log₁₀-transformed concentration, y = bottom + (top − bottom)/(1 + 10^((log EC50 − log[c]) × h)), where h is the Hill slope. Fitting was performed by non-linear least squares (trust-region reflective algorithm, scipy.optimize.curve_fit) on replicate-level data rather than on concentration means, so that each concentration contributed in proportion to the number of wells measured. Both asymptotes were fitted freely rather than constrained. Initial estimates for the upper and lower asymptotes were taken from the observed response range, and for log EC50 from the tested concentration whose mean response lay closest to half-maximal. The Hill slope was constrained to [0.05, 10] and log EC50 to the tested concentration range extended by one log-range in each direction. EC50 values are reported as 10^(log EC50). Goodness of fit was assessed by R². Fits whose EC50 fell outside the tested concentration range, or whose log EC50 was poorly constrained by the data (standard error > 0.5), were flagged and interpreted accordingly. For display, each curve was scaled to the mean response across its two highest concentrations (×100); no baseline was subtracted, and this scaling was applied after fitting, so reported EC50 values are derived from unscaled data and are unaffected. Analyses were performed in Python 3.11.3 using NumPy 2.4.6, pandas 3.0.3, SciPy 1.17.1 and Matplotlib 3.11.0.

### Cytotoxicity Assay

HLA-A*02:01 expressing K562 Antigen Presenting Cells were engineered to express GFP so that cell killing could be tracked by disappearance of fluorescence during a cytotoxicity assay. After pulsing with MART1-peptide for 2 hours, APCs were co-cultured with expanded T cell lines F12, A11, or H11 in microwells, and GFP fluorescence as well as IFN-γ secretion were monitored over time. 500 microwells loaded with at least 1 MART1 or DMSO-pulsed APC, and co-cultured with or without F12 T cells were analyzed and the disappearance of GFP scored as readout for cell death. Cell counts were derived from image segmentation and manually verified for every microwell. 95% confidence intervals were computed as Wilson score intervals for a binomial proportion using the Python package statsmodels version 0.14.6.

### TCR sequencing

QIAGEN RNAeasy Micro/Mini Plus Kit was used to extract RNA. DriverMap AIR TCR CDR3 (Cellecta, Inc., Cat: DMAIR2-HTR) was used to perform TCR sequencing. The quality of sequencing reads was assessed using FastQC^35^. Reads were then aligned with MiXCR^36^ using the DriverMap AIR preset. The TRA and TRB CDR3 sequences of each clone were obtained from the MiXCR output. Similarity of EC50 values between cell lines sharing a TCR was assessed by an exact permutation test. Group labels were treated as exchangeable across cell lines under the null hypothesis, and the test statistic was the mean absolute within-pair difference in log₁₀ EC50. The complete reference distribution was enumerated over all 3,150 distinct assignments of three disjoint pairs among the ten cell lines, giving an exact one-sided p-value. A rank-transformed version of the same test was run as a sensitivity analysis. Analyses were performed in Python 3.11.3 using NumPy 2.4.6 and pandas 3.0.3.

### TCR candidate expression in Jurkat-reporter cells

Each MART1-specific TCR sequence (assembled in structure as TRBV-TRBC-P2A-TRAV-TRAC ) was cloned in Cellecta’s cDNA expression vector pRR-EF1-MCS-Ubi-puro. Sequence verified TCR expression constructs pRR-EF1-TCR-Ubi-puro were packaged in 293FT cells using Cellecta’s packaging mix in 10 cm plates. Virus supernatants were collected 48hr later and concentrated by centrifugation at 18,500 rpm, 4°C for 2 hrs. The virus pellets were dissolved in PBS. The concentrated lentivirus expressing MART1-specific TCR were used to transduce Cellecta’s Jurkat NFAT-GFP reporter cells (Cat# ZJUR-T-Ble). Three days post transduction, the TCR expressing Jurkat reporter cells were selected by incubation in medium with 1 μg/mL puromycin for 4 days before expanding and maintaining in regular medium. To test the activation of each specific MART1 TCR construct, K562 cells that express Single-Chain-Trimer (SCT) MART1 (or NY-ESO1 as control) were used to coculture with above puro-selected TCR expressing Jurkat reporter cells. 1 × 10^5 Jurkat reporter cells (expressing TCR) and 0.4 × 10^5 K562 cells (expressing SCT_mart1, or SCT_NY-ESO-1, or without SCT) were plated in each well of 96well plate. After 24 hours of incubation, cells were collected and analyzed by flow cytometry for percentage of GFP+ cells.

### TCR-Antigen Pair Identification

A cell system consisting of two engineered cell lines was used to detect the antigen specificity of the TCR: (1) K562 APC line (Cellecta, Inc., Cat: YK562-MH-A0201-P) transduced with a lentiviral pool of 100 cancer antigens (Cellecta, Inc., Cat: CPLVPL-P) and (2) Jurkat TCR reporter cell line expressing a human CMV pp65 antigen-specific TCR (Cellecta, Inc., Cat: YJUR-TCMV-PBle). The cell lines were co-incubated in the cloneXplorer instrument for 24 hours. Candidate K562 APCs that were picked (along with the control transduced K562 cells) were processed for high-throughput sequencing with multiplex PCR using the DriverMap EXP assay protocol (Cellecta, Inc.) with custom primers specific to the 100 cancer antigens. The libraries were sequenced using NextSeq 2000 (Illumina, Inc.). The sequencing reads were aligned to the reference 100 cancer antigens using *Salmon*^37^.

### Cytokine Analysis

Celldom plates were primed with a solution containing 10 μg/mL IFN-γ capture antibody (clone NIB42 for human IFN-γ, clone R4-6A2 for murine IFN-γ, both BioLegend) to deposit the antibodies on the microwell array surfaces. Plates were incubated for 1-24 hours at 4°C, followed by backfilling with a non-adhesion coating (Anti-Adherence Rinsing Solution, Stemcell Technologies) to reduce cell migration. Soluble fluorescent IFN-γ detection antibody (PE-labeled clone 4S.B3 for human, clone XMG1.2 for murine, both BioLegend) was added to the assay medium so that the signal from individual microwells could be monitored continuously for more than 48 hours. We defined the relative secretion signal at timepoint t_x_ as the difference in the mean fluorescent intensity of the walls at timepoint t_x_ relative to the fluorescent signal at the initial timepoint t_0_. To eliminate false positives, we also quantified the rotational symmetry in the fluorescent pattern on the microwell walls, in which we reasoned that microwells with strong asymmetry are caused by cytokines captured from an adjacent microwell containing IFN-γ secreting cells. By gating the microwell parameter based on intensity and symmetry, we identified the true positives for cell retrieval. A second method for identifying microwells with IFN-γ secretion was developed by training a Faster-RCNN object detection model to find the instances of secretion rings directly. By using the intersection of AI detection methods with statistical methods involving intensity and symmetry measurements, we confidently identified the true positives.

### Mouse Model

OT-1 transgenic mice (C57BL/6 background) and wild-type C57BL/6J mice (6-8 weeks old, females) were purchased from the Jackson laboratory. All animal procedures were approved by MD Anderson Cancer Center Animal Care and Use Committee. The OT-1 transgenic mouse model expresses a T cell receptor specific for the OVA257–264 peptide (SIINFEKL) for the tracking and study of antigen-specific CD8+ T cell responses. We generated MC38 cells expressing ovalbumin via lentiviral transduction. OT-1 T cells were isolated from the splenocytes of OT-1 mice and expanded with Dynabeads Mouse T-activator CD3/CD28 (ThermoFisher) before loading on Celldom plates. Antigen-presenting cells were differentiated from the bone marrow of mice.

## Acknowledgements

The authors are grateful for the support of this research by National Institutes of Health (R44CA281575, R35GM122465, DL119795). X.S. is a Cancer Prevention and Research Institute of Texas (CPRIT) Scholar in Cancer Research and this work was supported under grant (RP240007).

## Author Contributions

G.S. designed and performed the biological experiments on the cloneXplorer. B.B.Y developed the image analysis pipeline, including the training and deployment of the AI models, and the user interfaces for the cloneXplorer and Distiller. M.Z. developed the firmware and synchronization for the instrument. O.N. and E. T. designed and fabricated the microwell plates. O.Z. supervised the training and deployment of computer vision models. S.S, H.R., and X.S. established the mouse models. D.D., K.P., and A.C. developed the K562 and Jurkat models and performed next generation sequencing and data analysis. All authors contributed to editing of the manuscript, discussion, and development of figures, and all authors approve this manuscript.

## Data availability

Data supporting the findings of this study are available within the article, its supplementary files, and in the Bioimage Archive^12^. The sequencing data generated in this study have been deposited in the NCBI Gene Expression Omnibus (GEO) under accession number GSE326895. Any additional requests for information can be directed to, and will be fulfilled by, the corresponding authors. Source data is provided with this paper.

## Code availability

Custom code developed by Celldom, Inc. was used to interface with the instrument, acquire and analyze the data. A custom data pipeline developed by Celldom, Inc. was used to output cell counts and secretion measurements from the raw electronic data. All custom code is proprietary to commercial products and is not publicly available.

## Abbreviations

APC: Antigen Presenting Cell
BF: brightfield
BMDC: Bone Marrow Derived Dendritic Cell
CDR3: Complementarity-Determining Region 3
DC: Dendritic Cell
GFP: Green Fluorescent Protein
IFN-γ: Interferon-gamma
MART1: Melanoma Antigen Recognized by T Cells 1
NFAT: Nuclear Factor of Activated T cells
OME-NGFF: Open Microscopy Environment Next Generation File Format
PBMC: Peripheral Blood Mononuclear Cells
SCT: Single Chain Trimer
scTCR-seq: Single Cell TCR sequencing
TCR: T Cell receptor

## Supplementary Analysis of Co-Culture Interaction Combinations in Microwells

We aim to estimate the maximum number of TCR/epitope combinations that can be screened in one experiment. Let **N** be the number of microwells filled with an average occupancy, **Λ**, cells per microwell, such that **M = N Λ** is the total number of cells detected in the observed microwells. The total cell input consists of two populations: **N Λ = N (Λ_1_ + Λ_2_**), where **Λ_1_** is the average occupancy of the TCR-expressing Jurkat cells and **Λ_2_** is the average occupancy of the antigen-presenting K562 cells (APCs). Assuming each individual **J** species in the TCR-expressing population is equally represented as **λ_j_ = Λ_1_ / J** for j∈{1 .. J} and similarly for the **K** species of APCs with probabilities as **λ_k_ = Λ_2_ / K** for k∈{1 .. K}, then the joint probability of finding a j/k pair in any given microwell is **λ_jk_ = (Λ_1_ / J) (Λ_2_ / K).** The ability to observe each j/k pair requires enough microwells to satisfy: **α N λ_jk_ >> 1**, where **α**∈[0,1] is the probability that a positive signal is observed in a microwell that contains a matching TCR/antigen pair and is ideally close to unity. In this work, we found that **α** ∼ 0.7, meaning that GFP+ microwells are observed in 70% of instances where there is a matching TCR/antigen pair. With substitution, we can estimate the upper limit of the assay as: **α N (Λ_1_ / J)(Λ_2_ / K) >> 1**.

To evaluate scaling trends, assume **J ∼ K** and **Λ_1_ ∼ Λ_2_**, implying an equal number of Jurkat and K562 cells are loaded into the microwells, and their libraries have equal plexity. This yields **P < Λ**^2^ **(α N / 4),** where **P** is the maximum plexity for given values of **Λ** and **N**. This relationship shows that the optimal plexity increases linearly with the number of cells loaded, which makes sense because the number of interactions occurring in microwells scales quadratically with the number of cells inside. The plexity depends linearly on the number of microwells. For example, if there are **N = 270,000** pickable microwells and with **Λ ∼ 2** and **α ∼ 1**, a library of size ∼165×165 will yield ∼10 co-culture pairs for each J/K combination. Since the recovered sequences are derived from small cell populations of ∼ 2 – 5 cells, the correctly matched j/k pairs must occur at sufficient frequency to offset counts from bystander sequences recovered from those microwells. To validate the interaction, the j/k pairs must be observed in the sequences recovered from at least 3 independent microwells.

This analysis suggests the possibility of screening millions of TCR/antigen pairs simultaneously by loading the microwells with dozens of cells per well. However, this throughput can only be achieved by reducing background noise caused by randomly activated T cells, which pollute the deconvolution pipeline. Background noise can be reduced by pre-sorting the Jurkat population with a GFP-negative gate, or better yet, by identifying the pre-activated T cells during live cell imaging. The ability to image microwells at multiple time points allows false positives to be eliminated, which is an advantage over systems that rely on single time point imaging or endpoint genomic assays. If background can be reduced below 0.001% using a gating scheme based on multiple timepoints and functional parameters, then it should be possible to screen 100,000 TCR/epitope interactions in one plate. This platform can handle both single library screens, such as one TCR evaluated against a pool of 100,000s of antigens, as well as library-on-library screens, such as the validation of a short list of 100 TCR candidates against a smaller library of 1,000 antigens.

---

**Supplementary Fig. 1.**
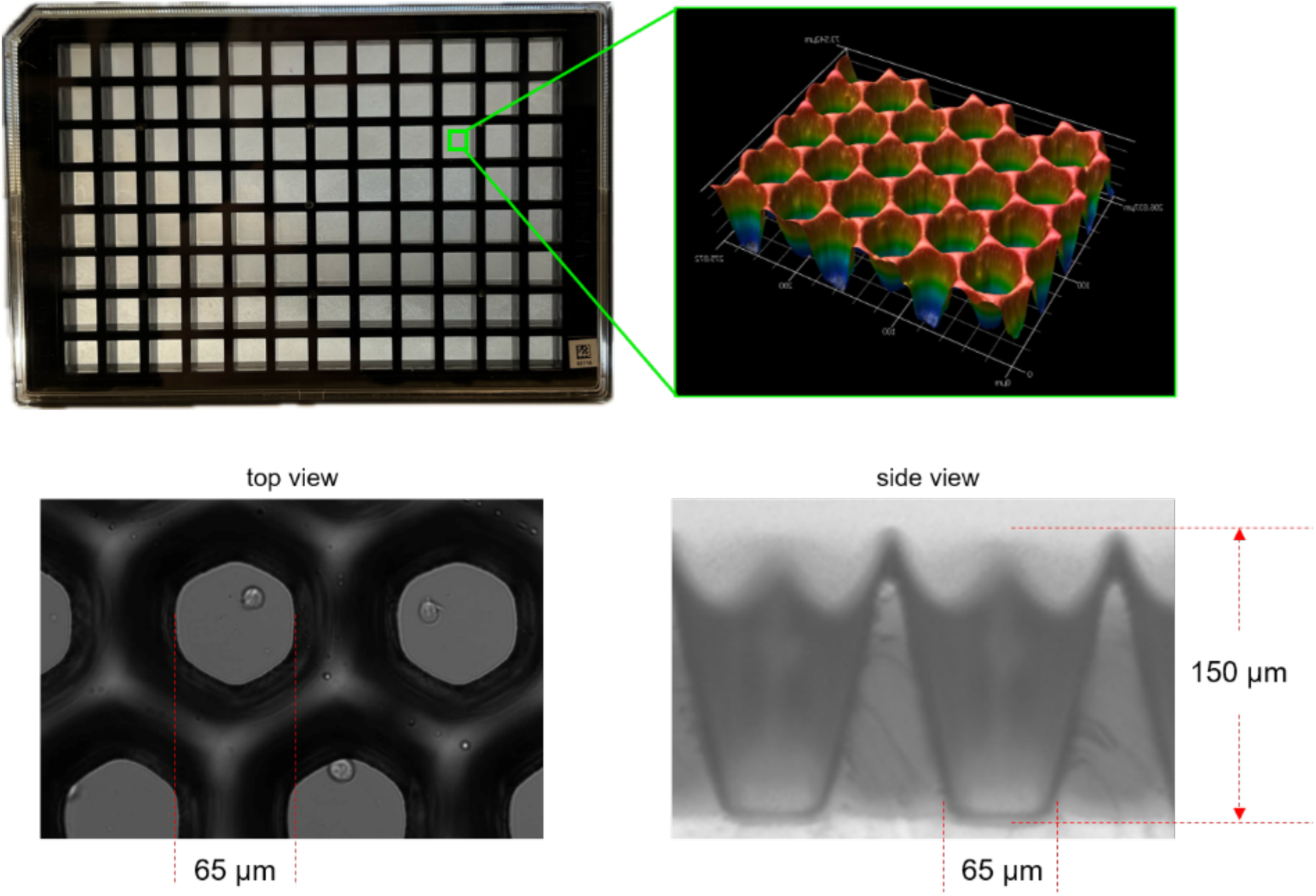
(Top Left) A photograph of the microwell plate. (Top Right) A magnified view of the microwells measured by optical profilometry. (Bottom Left) A top view of microwells loaded with cells. (Bottom Right) A side view of the microwells.

**Supplementary Fig. 2.**
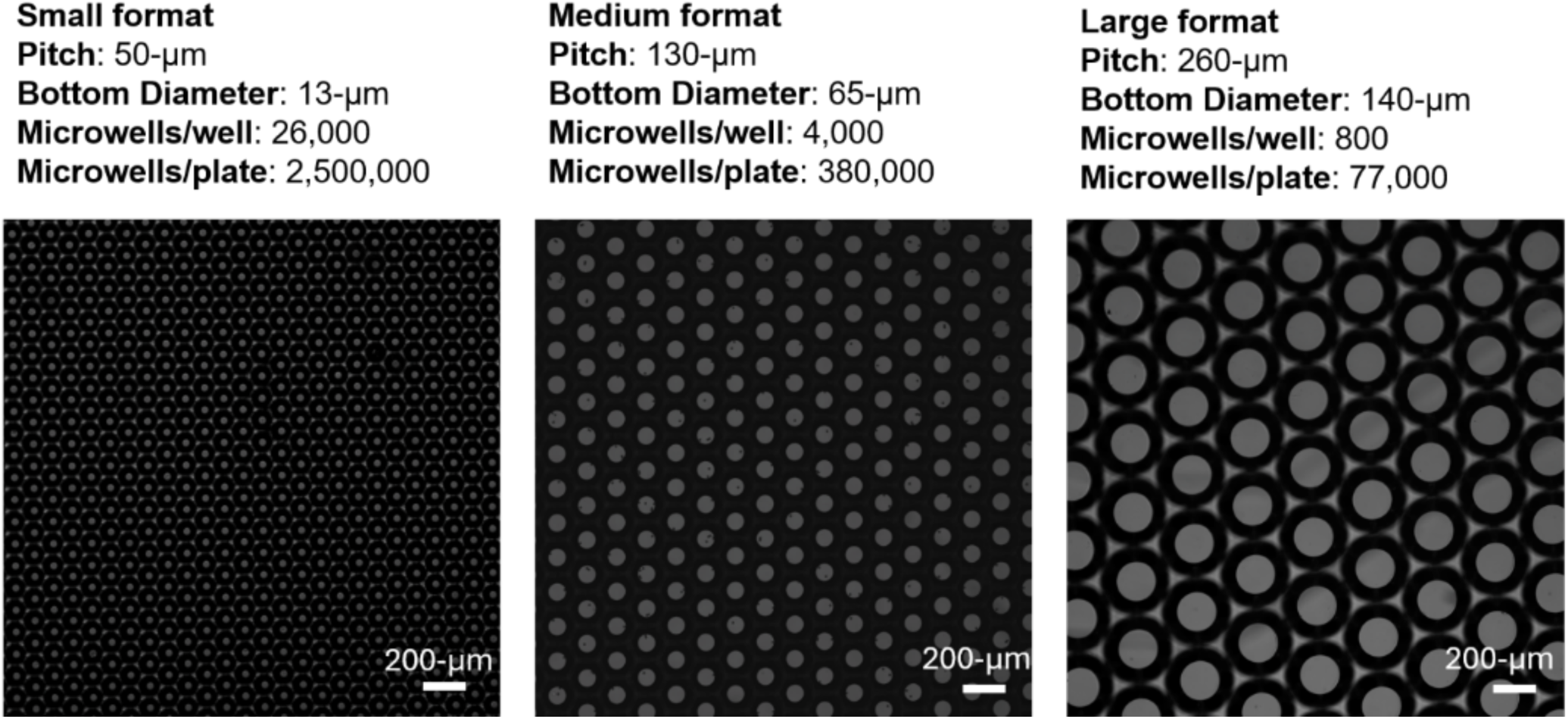
Comparison of microwells with different sizes that Celldom manufactures for different applications. The medium format is used in this work.

**Supplementary Fig. 3.**
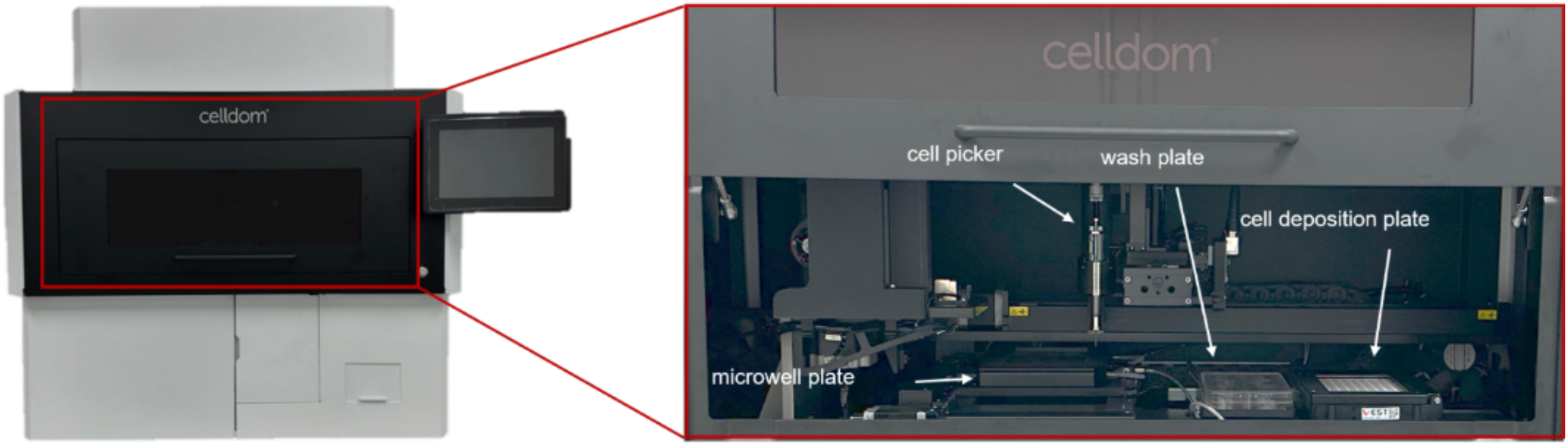
cloneXplorer instrument with closed door (left), and open door (right).

**Supplementary Fig. 4.**
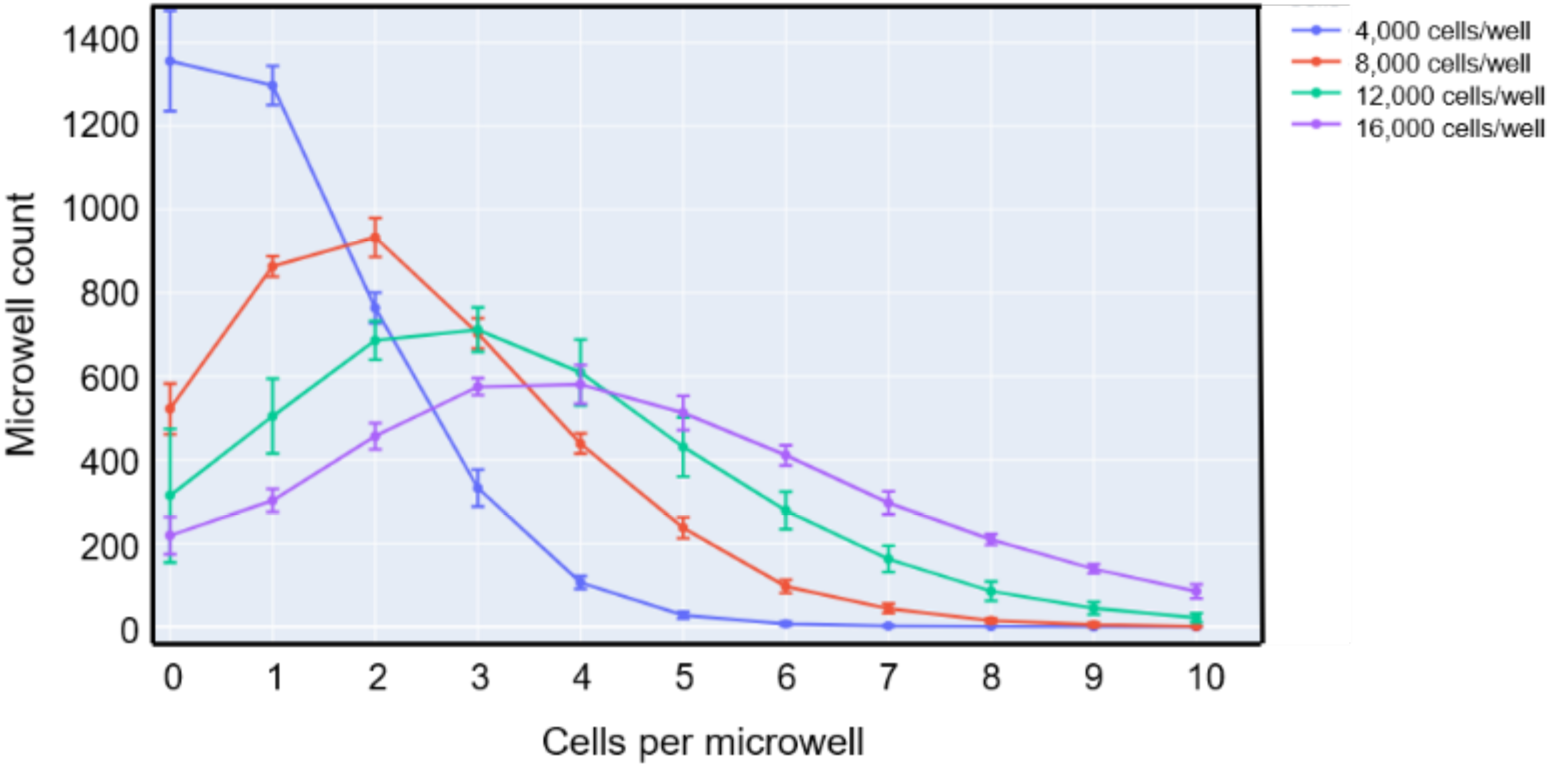
Cell loading distribution for 4,000, 8,000, 12,000, and 16,000 cells seeded per well. For each condition, mean and standard deviation of 12 wells are plotted.

**Supplementary Fig. 5.**
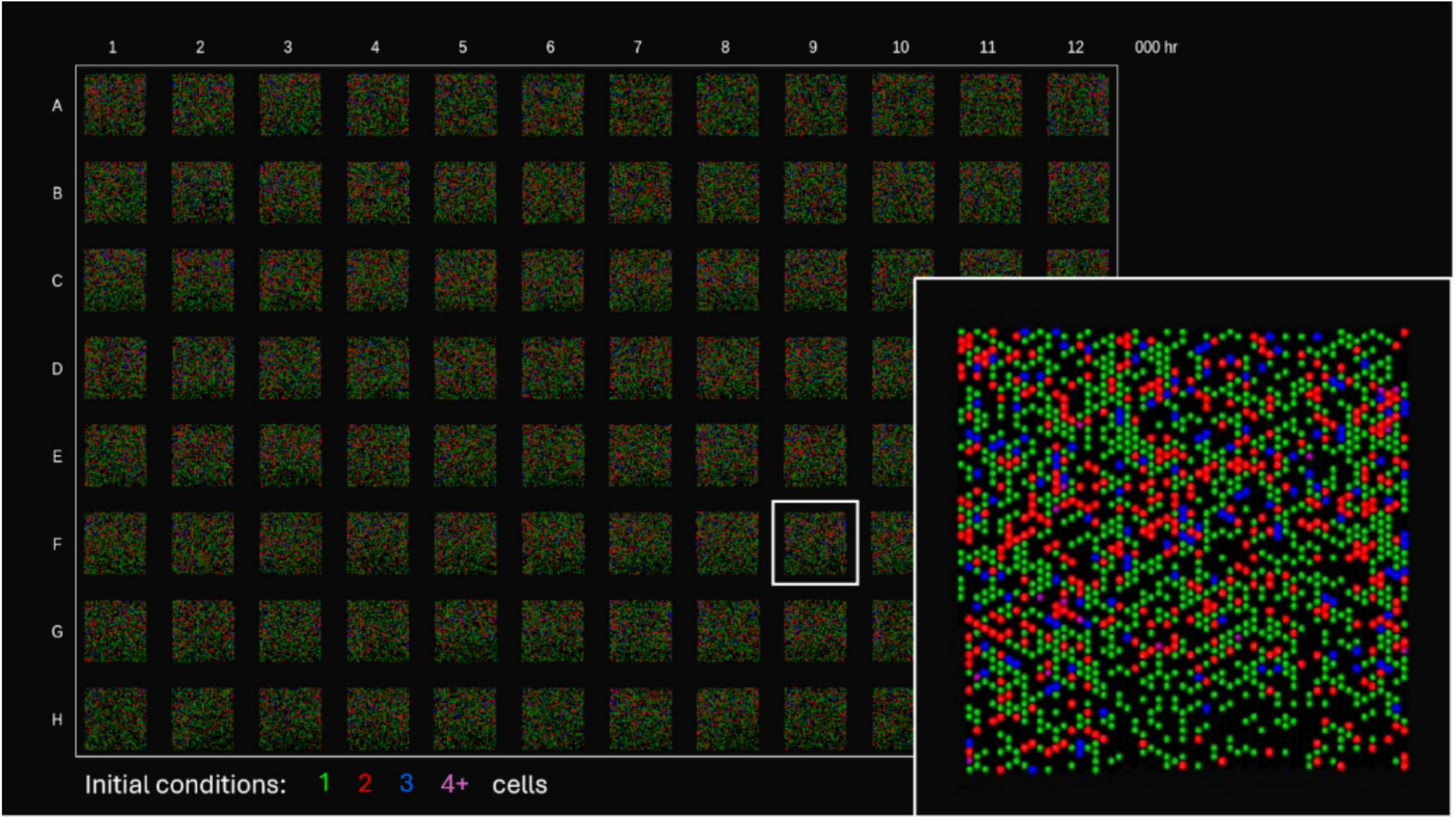
Cell loading distribution for 3,300 cells per well. The black, green, red, blue, and purple colors indicate microwells that contain 0, 1, 2, 3, or 4+ cells.

**Supplementary Fig. 6.**
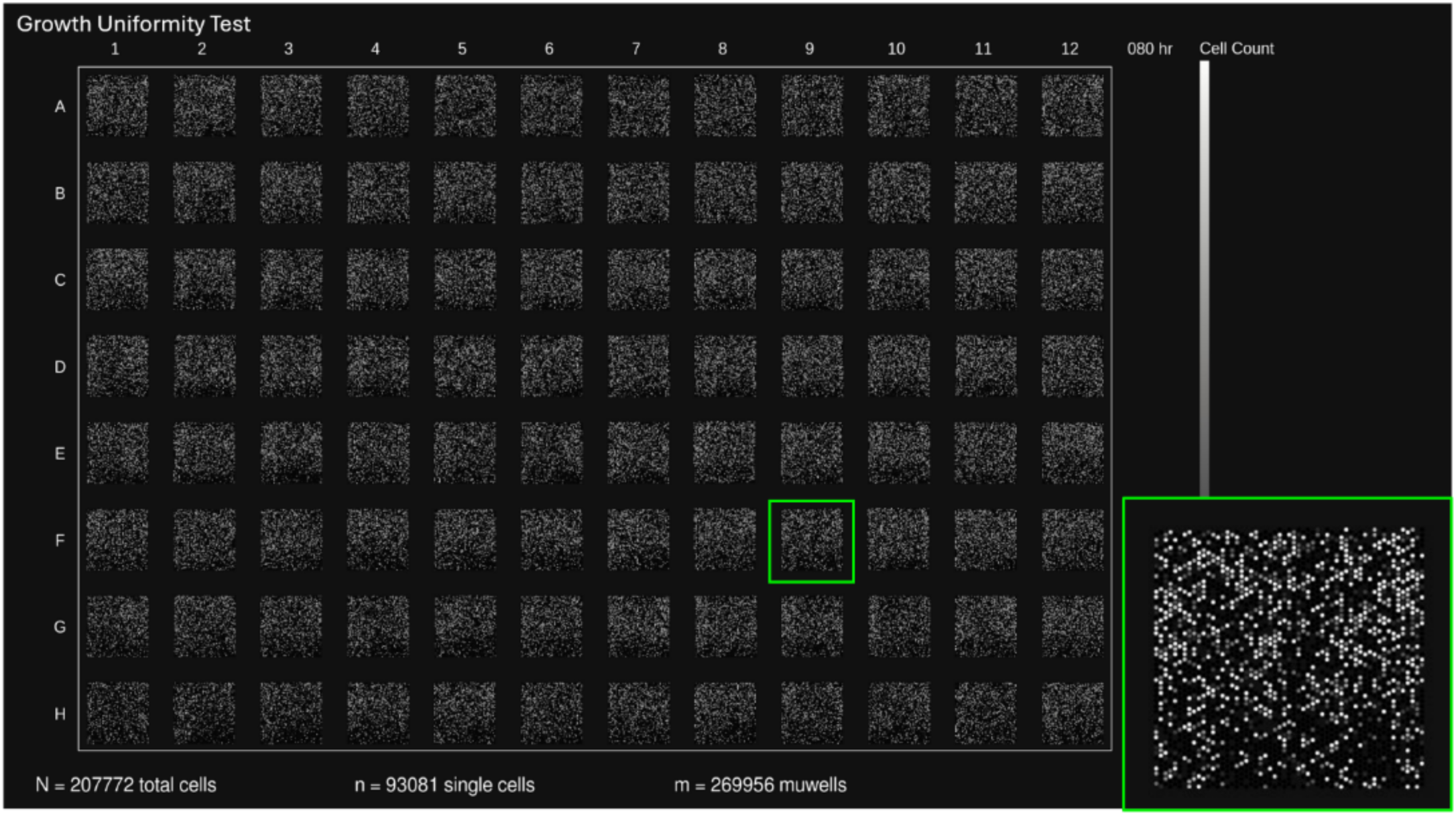
In the microwells that initially had a single cell, the cell count map is shown at 80 hours with grayness level proportional to the number of cells. The whitest color is 10 cells.

**Supplementary Fig. 7.**
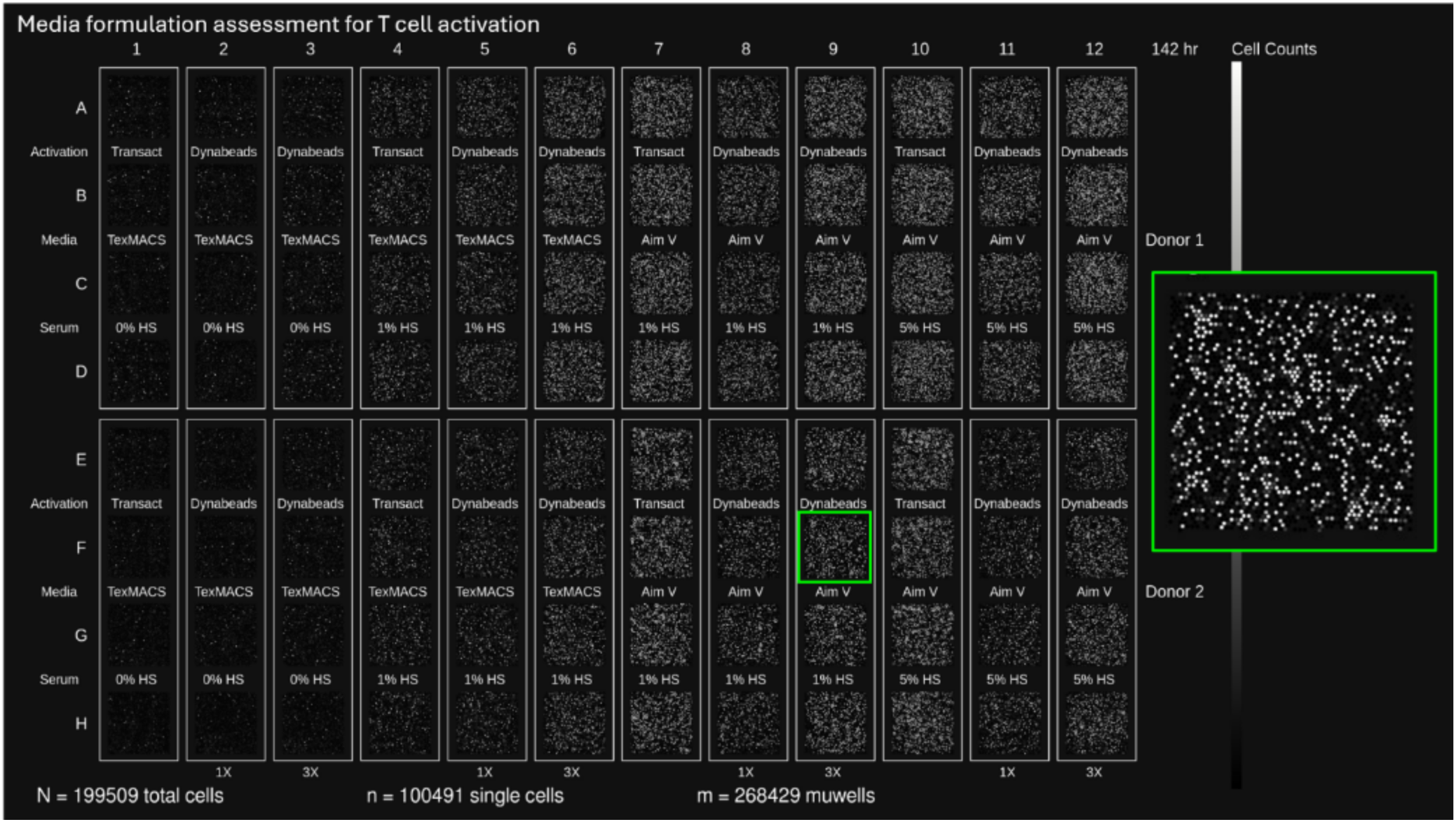
Cell growth distribution for freshly thawed T cells exposed to different levels of human serum (0%, 1%, 5%), different methods of CD3/CD28 stimulation (Dynabeads vs Transact), and different media conditions (TexMACS vs AIM V). The gray level indicates the cell count at 142 hr on a scale of 0 (black) to 100 (white).

**Supplementary Fig. 8.**
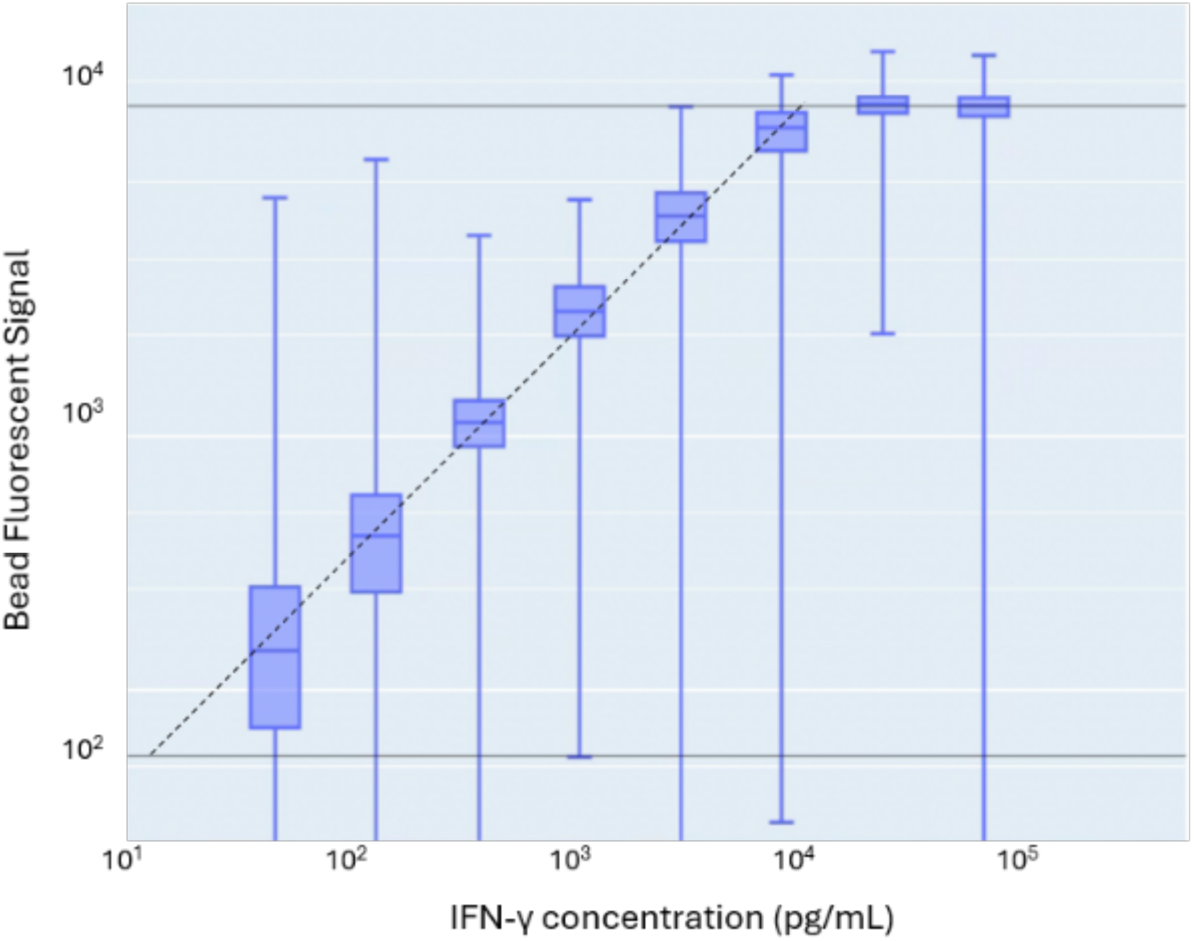
Fluorescent signal from IFN-γ capture beads in microwells exposed to different concentrations of soluble cytokine IFN-γ.

**Supplementary Fig. 9.**
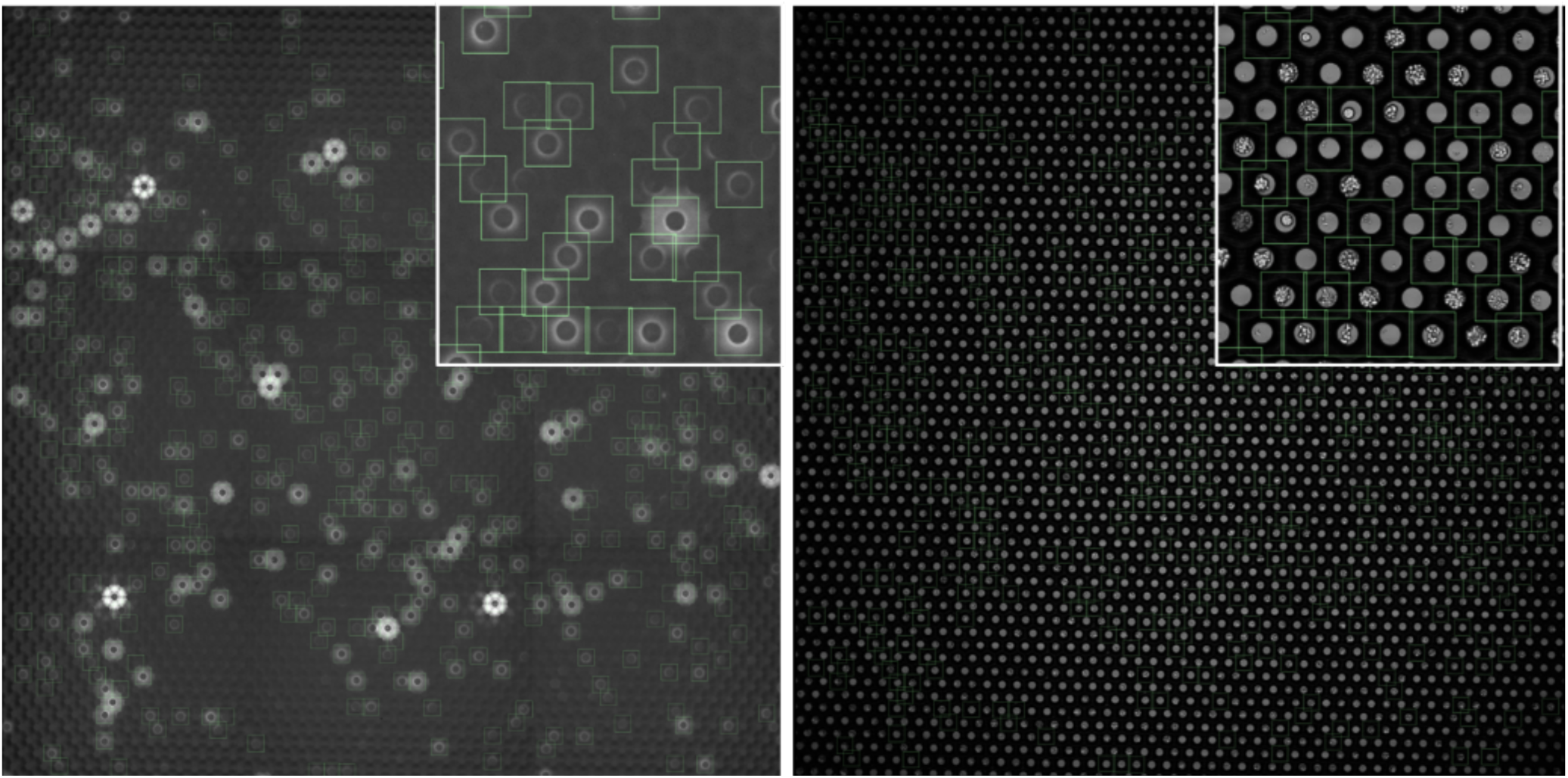
Whole well stitch of IFN-γ secretion pattern (left) and the corresponding brightfield images (right) in a coculture of mouse OT-1 cells and MC38 cells presenting SIINFEKL peptide.

**Supplementary Fig. 10.**
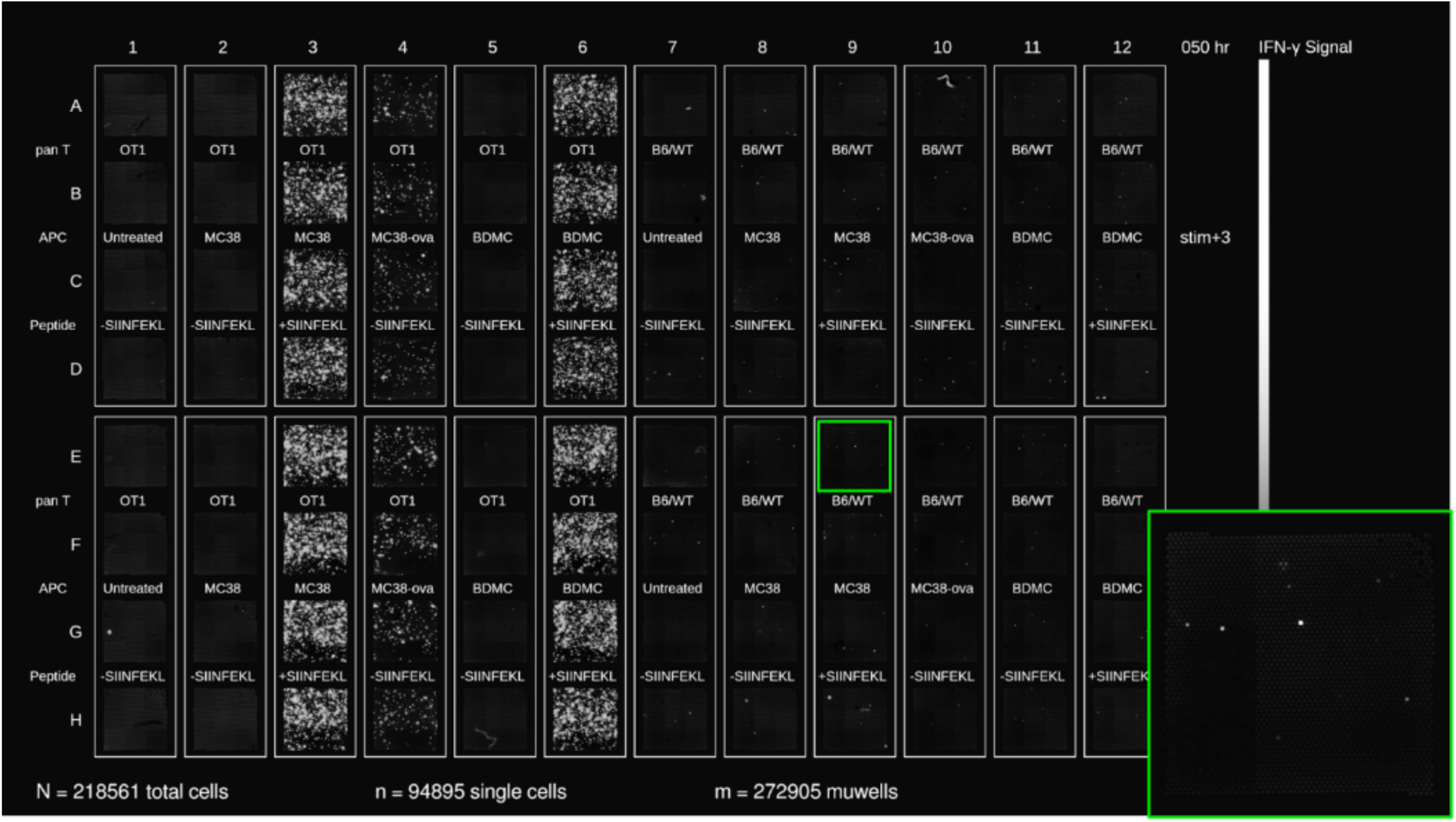
A microwell map of IFN-γ secretion at 50 hours is shown for co-cultures of splenocyte derived mouse T cells (OT-1 vs B6/WT) in co-culture with different antigen presenting cells (MC38, MC38-ova, BMDC) in the presence or absence of SIINFEKL peptide.

## SUPPLEMENTARY MOVIE LEGENDS

**MOVIE S1.** 7 μm fluorescent beads are retrieved from microwells in 250 successive picks.

**MOVIE S2.** NALM6-GFP cells are retrieved from microwells in 35 successive picks.

**MOVIE S3.** A movie of a single K562 cell imaged at 4 hour intervals for 56 hours is shown with and without the segmentation masks.

**MOVIE S4.** To assess growth rate uniformity across a 96-well plate, the NALM6 cell count is shown at 8 hour intervals for 80 hours. Out of the 207,772 cells identified at the initial timepoint, 93,081 were single cells.

**MOVIE S5.** To assess uniformity in the proliferation rates of primary T cells, freshly thawed T cells were exposed to Miltenyi Transact in AIM V media supplemented with 5% human serum and imaged at 3 hour intervals for 130 hours.

**MOVIE S6.** To assess uniformity in the proliferation rates of primary T cells, freshly thawed T cells were exposed to ThermoFisher Dynabeads in AIM V media supplemented with 5% human serum and imaged at 3 hour intervals for 130 hours.

**MOVIE S7.** The activation dynamics of freshly thawed human T cells exposed to different CD3/CD28 activation matrices (Miltenyi Transact vs ThermoFisher Dynabeads) in different media (AIM V vs TexMACS) and supplemented with different levels of human serum (0%, 1%, 5%) is shown as a map of cell counts imaged at 3 hour intervals for 130 hours. Out of the 199,509 cells identified at the initial timepoint, 100,491 were single cells.

**MOVIE S8.** The life of a single cell from the day of seeding through 3 days of expansion is shown, followed by documentation of the cell picking process, including images taken before and after cell isolation in brightfield and fluorescent channels.

**MOVIE S9.** Time lapse brightfield images are shown for co-cultures of mouse T cells (OT-1) and MC38 ova cells.

**MOVIE S10.** The characteristic IFN-γ secretion patterns are shown for co-cultures of mouse T cells (OT-1) and MC38 ova cells at 4-hour intervals for 50 hours.

**MOVIE S11.** A secretion map of mouse T cells (OT1 vs B6/WT) in co-culture with antigen presenting cells (MC38, MC38-ova, BMDC) and exposed to SIINFEKL peptide is shown at 4 hour intervals for 50 hours. Out of the 218,561 cells identified at the initial timepoint, 94,895 were single cells.

**MOVIE S12.** A single T cell from cell line F12 kills 3 GFP-labeled MART1-pulsed K562 APCs. Full movie of which selected time points are shown in Fig. 6i.

**MOVIE S13.** A single T cell from cell line F12 is overgrown by GFP-labeled DMSO-pulsed K562 APCs. Full movie of which selected time points are shown in Fig. 6j.

